# A response-geometry framework separates microbiome movement magnitude from directional coherence in intervention studies

**DOI:** 10.64898/2026.05.22.726133

**Authors:** Carol Ying Ying Szeto, Hoi Shan Kwan

## Abstract

Dietary and lifestyle microbiome interventions often produce mild but heterogeneous remodeling rather than uniform community shifts. In this setting, scalar diversity or group-level summaries can appear weak or inconclusive even when participants move in organized but magnitude-limited directions, or move substantially in divergent directions. We developed a response-geometry framework that jointly describes baseline-referenced response magnitude and cross-participant directional coherence within a compositional feature space. The framework complements diversity, ordination, trajectory, PERMANOVA, PERMDISP, and beta-diversity analyses by asking whether paired responses differ in size, shared direction, or both.

**Methods:** A response vector for each participant was defined as the follow-up minus baseline profile after adding a 0.5 pseudocount and applying centered log-ratio transformation in Aitchison-based response space. Response magnitude was the Euclidean length of this vector. Directional coherence was quantified as cosine alignment between participant-level response vectors and the mean group response vector, with sign-flip permutations as a paired-structure-preserving diagnostic null. We evaluated the framework using workflow-sensitive diversity comparisons, 198,000 logistic-normal compositional simulations with 100 or 500 features and small-to-large shared-direction effects, public-data-derived implementation stress tests, a synbiotic and dietary-intervention cohort, and a fiber/fermented-food application in 16S rRNA gene amplicon and shotgun-derived CAZyme gene-family feature spaces. A beta research-preview repository accompanying the preprint is available at https://github.com/carolyyszeto/microbiome-response-interpreter-beta as v6.5-beta, including documented scripts, a toy dataset, environment notes, output-interpretation guidance, and exploratory implementation utilities.

**Results:** Workflow comparisons showed that richness-sensitive differences were concentrated in rare-tail and low-abundance structure, informing the analytical feature-space context for response interpretation. In simulations, null and magnitude-only random-direction scenarios showed near-null detection rates of 0.061 and 0.062, close to nominal alpha = 0.05, whereas shared-direction scenarios showed increasing coherence with stronger effects and larger sample sizes. Mixed-responder and opposing-subgroup scenarios attenuated or cancelled pooled coherence, supporting separation between response magnitude and directional organization. The synbiotic and dietary-intervention cohort showed modest, heterogeneous displacement with limited within-arm coherence, with permutation p values from 0.575 to 0.653. In the fiber/fermented-food application, fermented-food exposure showed stronger 16S response organization than the baseline-period reference, while CAZyme estimates used non-identical sampling endpoints and remained feature-space-specific.

**Conclusions:** This response-geometry framework helps distinguish paired microbiome movement size from shared response orientation. It is intended as an interpretively cautious response-organization descriptor for mild, heterogeneous intervention settings, not as a replacement for existing multivariate methods. Its interpretation depends on sample size, effect structure, endpoint alignment, zero handling, group-direction stability, and feature-space definition. The framework does not convert weak, null, endpoint-limited, or sensitivity-dependent findings into efficacy, predictive, or mechanistic claims.

## Background

Dietary and lifestyle microbiome interventions often produce limited and heterogeneous remodeling rather than uniform community shifts, with diet-microbiome associations varying across individuals and time [1]. In this setting, a weak or inconclusive scalar diversity result does not necessarily mean absence of microbiome response, as diet-targeted interventions can show layer-specific taxonomic, functional, or host-context patterns [2]. A large overall displacement also does not necessarily mean that participants moved in a shared ecological direction, because dispersion or distance from a reference can increase without specifying whether response vectors share orientation [3, 4]. Participants may instead show modest but organized movement, substantial but divergent movement, or feature-space-dependent changes, a distinction consistent with trajectory-based community analysis in which path length and direction can represent different aspects of temporal community change [5]. This interpretive problem is especially relevant for dietary, synbiotic, probiotic, prebiotic, fiber, fermented-food, and lifestyle-style interventions, where responses can be personalized or baseline-dependent in probiotic, synbiotic, fiber, and prebiotic contexts [6, 7, 8, 9].

Microbiome diversity remains a useful compact summary of community state and intervention-associated change. Alpha-diversity measures summarize within-sample richness and evenness, whereas beta-diversity distances and ordination approaches describe between-sample community differences [10, 11, 12, 13, 14]. These summaries reduce high-dimensional microbial community profiles into interpretable quantities, but they do not by themselves explain why a community appears to have changed. Scalar diversity change captures within-sample richness and evenness but compresses community structure into one value [15, 14]. Total displacement or beta-distance summaries quantify between-sample separation in ecological or phylogenetic distance space [11, 12]. Group-centroid methods such as PERMANOVA evaluate location-like differences among groups [16], whereas dispersion summaries such as PERMDISP and distance-to-centroid beta-diversity describe multivariate spread [17, 3]. Each summary can capture important aspects of microbiome remodeling, yet each can also collapse distinct paired-response properties.

Analytical context further complicates this interpretation. Sequencing workflow, feature definition, taxonomic assignment, abundance filtering, sampling depth, and metric choice can shape which taxa enter the community table and how community change is represented [13, 14, 18]. Normalization and rarefaction choices influence diversity estimation, and no normalization choice is universally optimal across analysis goals and diversity questions [19, 15, 20, 21]. Compositional constraints and differential-abundance method choice can materially alter microbiome results, especially when relative-abundance data are interpreted without accounting for compositional structure [22, 23, 24, 25]. This issue is especially important for richness-oriented metrics, where rare-tail and low-abundance features can drive apparent diversity shifts. Low-abundance taxa may carry ecological information, but they can also be unstable, detection-sensitive, or artifact-prone in amplicon datasets [26, 27, 28]. Analytical choices can therefore affect both the scalar readout and the feature space in which response magnitude and direction are later assessed.

Longitudinal intervention studies introduce an additional paired-response problem after the analytical space has been defined, because repeated-measures microbiome data require explicit representation of within-participant change over time [29, 30]. Participants may move different distances from baseline, a concept related to trajectory length or displacement in multivariate community space [5]. They may or may not move in a shared ecological direction, because movement orientation is a directional property that is distinct from movement size [4, 5]. A shared direction means that participants show similar multivariate shifts in community composition within the same feature space, rather than merely similar changes in a single diversity value. Existing methods provide important but different information. Alpha-diversity change summarizes within-sample scalar change in richness or evenness [15, 14]. Beta-diversity displacement summarizes between-sample separation in non-phylogenetic ecological distance space [11]. Phylogenetic beta-diversity frameworks provide complementary distance-based community comparisons that incorporate evolutionary relatedness among taxa [12]. PERMANOVA and PERMDISP test group-location and dispersion structure, with multivariate dispersion also used as a beta-diversity descriptor based on distances to group centroids [16, 17, 3]. Compositional ordination and perturbation-oriented approaches help represent microbiome change in log-ratio or compositional feature space [31, 22, 32]. Time-dependent community methods, including principal response curves and temporal beta-diversity frameworks, provide established approaches for analysing multivariate community change over time [33, 34]. Longitudinal microbiome workflows and log-ratio approaches further help model repeated-measures structure and compositional change over time [29, 35, 30]. These approaches do not directly separate how far each participant moved from their own baseline from whether participant-level response vectors were aligned across participants.

This separation matters because mild or heterogeneous intervention responses can be misread when response size and response orientation are not distinguished. Participants may show comparable displacement magnitudes but move in divergent directions, illustrating that movement size and movement orientation are separable descriptors in multivariate space [4]. Participants may also show modest displacement while still moving along a shared response direction, a distinction consistent with trajectory-based community analysis in which path length and direction describe different aspects of temporal community change [5]. However, microbiome intervention studies have not routinely applied baseline-referenced displacement and cross-participant directional alignment together as paired descriptors that jointly characterize response magnitude and directional organization.

We develop a computational response-geometry framework for paired microbiome remodeling that formalizes this distinction as a complementary layer to existing methods. The framework separates response magnitude, defined as how far each microbiome profile moves from its own baseline, from directional coherence, defined as whether participant-level response vectors are aligned across participants within the same analytical feature space. Directional coherence is quantified using cosine similarity. The framework is intended as a cautious response-organization descriptor that helps distinguish organized response, heterogeneous movement, magnitude-only displacement, and endpoint-caveated functional-context patterns without converting weak, null, or sensitivity-dependent findings into efficacy or mechanistic claims.

The manuscript applies this framework across analytical and empirical contexts. Workflow-sensitive diversity comparisons examine how richness and rare-tail structure shape the feature-space context for response interpretation. An r500 simulation benchmark defines operating behavior under null, magnitude-only, shared-direction, mixed-responder, opposing-subgroup, and rare-tail-sensitive scenarios. David and Palleja implementation stress tests provide empirical operating contrasts, with the David animal-diet dataset used as a magnitude-stress contrast and the Palleja antibiotic-recovery dataset used as a strong-perturbation contrast across species and pathway feature spaces [36, 37]. Public dietary and synbiotic intervention datasets then provide practical application support. The Lauw synbiotic and dietary-intervention cohort is used as a modest, heterogeneous intervention-period worked example [38], and the FeFiFo/Wastyk fiber and fermented-food dataset is used to examine response organization in 16S rRNA gene amplicon taxonomic space and endpoint-caveated shotgun-derived CAZyme gene-family feature space [2].

## Materials and Methods

### Study design and analytical scope

This study evaluated microbiome change using a response-geometry framework designed to complement scalar diversity and conventional beta-diversity summaries. Workflow-comparison analyses examined how sequencing and processing contexts affected richness, diversity, and rare-tail structure. These analyses defined the analytical feature-space context in which later response-vector summaries were interpreted. An r500 simulation benchmark and supplementary implementation stress tests then examined the operating behavior of the directional-coherence descriptor before its application to modest intervention data. The framework was subsequently applied to a longitudinal synbiotic and dietary-intervention worked example and to an additional public dietary-intervention dataset. The workflow-comparison analyses examined how analytical sensitivity affected diversity readouts, richness geometry, and rare-tail structure across sequencing platforms and processing pipelines. The intervention analyses then represented each participant’s microbiome change as a paired baseline-referenced response vector and summarized response organization using response magnitude and directional coherence.

The longitudinal synbiotic and dietary-intervention cohort was used as a modest-perturbation worked example for evaluating microbiome response magnitude and directional coherence [38]. The r500 simulation benchmark examined the expected behavior and operating boundaries of the coherence descriptor under controlled response scenarios. Public-data-derived David/Palleja analytical outputs were used as supplementary implementation stress tests under stronger perturbation and magnitude-stress settings [36, 37]. The FeFiFo (Fermented and Fiber-rich Foods) dietary-intervention dataset extended the framework to 16S rRNA gene amplicon response space and to a constrained metagenomics-derived CAZyme gene-family response space [2].

Physiological, clinical, and functional-context analyses were handled as secondary or exploratory analyses. No formal power calculation was conducted because all intervention analyses were secondary reanalyses of fixed public datasets. No prospective analysis plan was preregistered. Statistical uncertainty was summarized using permutation, bootstrap, and sensitivity analyses. Endpoint-level analyses and supporting sensitivity outputs were reported in the corresponding supplementary tables rather than selected only when directionally positive.

### Data sources

Six public datasets or public-data-derived analysis outputs were used.

i. A public Illumina stool-donor cohort comprising 200 samples from 170 donations and 86 donors was obtained from the dataset reported by Santiago and Olesen [39] (ENA accession PRJEB41316). This cohort was used primarily for workflow-dependent comparisons of observed richness, Shannon index, Simpson index, and rare-tail behavior.
ii. A paired multi-platform human fecal dataset (n = 6) was based on the study by Matsuo et al. [40], in which the same six fecal samples were analyzed across nanopore and Illumina 16S sequencing contexts (DDBJ DRA accession DRR225043-DRR225065). This dataset was used for both cross-platform comparison and within-nanopore workflow comparison.
iii. A longitudinal intervention cohort was based on the randomized trial reported by Lauw et al. [38], comprising a synbiotic supplementation group (SG, n = 19), a dietary intervention group (DG, n = 18), and a combined dietary plus synbiotic group (DSG, n = 18). The original microbiome sequencing data were deposited under BioProject PRJNA1015382. This cohort served as a worked example for microbiome response interpretation in a modest, heterogeneous intervention setting.
iv. The Wastyk/FeFiFo fiber/fermented-food dietary-intervention dataset reported by Wastyk et al. [2] included high-fiber and high-fermented-food arms and was used as an additional public intervention application of the framework. The dataset contained processed longitudinal 16S rRNA gene amplicon profiles and metagenomics-derived CAZyme gene-family abundance profiles. FeFiFo paired sample sizes differed by data layer and endpoint contrast. The primary 16S rRNA gene amplicon endpoint retained 17 fiber-arm and 16 fermented-arm pairs, and the CAZyme gene-family endpoint retained 16 fiber-arm and 14 fermented-arm pairs. The full endpoint-specific sample-size summary is reported in Supplementary Table 10.
v. David 2014 public animal-diet data output was used as a supplementary magnitude-stress implementation contrast [36]. The available Animal-arm anchor transition was used to examine whether large displacement alone implied shared directional organization. This analysis was used as implementation stress-test support only.
vi. Palleja 2018 public antibiotic-recovery data outputs were used as supplementary strong-perturbation implementation contrasts [37]. Species and pathway feature spaces were analyzed separately to examine organized disruption and recovery-like directional behavior. These analyses were used as implementation stress-test support only.

### Illumina reprocessing and OTU-oriented comparison

Illumina raw sequence data were reprocessed within the present analytical framework. Taxonomic assignment across Illumina-based analyses used SILVA NR99 v138.1 as the taxonomic reference in the current pipeline [41, 42].

Paired-end reads were processed using DADA2 [43] for ASV inference. Forward and reverse reads were filtered and trimmed using filterAndTrim with working parameters truncLen = c(240, 200), maxN = 0, maxEE = c(2, 4), truncQ = 2, and rm.phix = TRUE. These parameters were selected after confirming that filtering and paired-read merging produced usable retained reads for reproducible reprocessing of the public dataset. They were retained as fixed reproducible settings rather than as universal DADA2 defaults. Error learning, dereplication, sample inference, paired-read merging, sequence-table construction, and chimera removal then followed the standard DADA2 workflow. Chimeras were removed using removeBimeraDenovo with the consensus method.

Representative ASV sequences were written to FASTA format and clustered at 97% sequence identity using VSEARCH with the --cluster_fast option to enable OTU-oriented comparison [44]. The 97% threshold was used as a conventional OTU-oriented comparison point rather than as a claim that OTUs are biologically preferable to ASVs. The resulting.uc assignments were used to construct an ASV-to-OTU mapping. OTU abundance matrices were generated by summing abundances of ASVs assigned to the same OTU cluster. OTU- and ASV-derived diversity metrics were then compared within harmonized sample sets.

### Diversity metrics, filtering, and rarefaction

Alpha diversity was summarized using observed richness, Shannon index, and Simpson index [45], with Chao1 included where relevant for the ASV-OTU inflation comparison. Observed richness was used primarily to assess taxon detection sensitivity and rare-tail expansion. Shannon and Simpson indices were used as abundance-weighted descriptors of community structure.

Beta diversity was assessed using Bray-Curtis dissimilarity [11] and Aitchison distance [31, 22]. Bray-Curtis was used for abundance-based ecological comparison. Aitchison distance was used for compositional paired-response analyses after centered log-ratio (CLR) transformation.

Common samples in matched ASV-OTU comparisons were filtered by sequencing depth before rarefaction. The matched PRJEB41316 ASV/OTU comparison included 159 matched samples before depth filtering. The working analysis scripts retained 148 samples after requiring both ASV and OTU depths of at least 5,000 reads, excluded 11 samples, and rarefied both workflows to 5,000 reads using seed 123. This threshold was used as a practical compromise between retaining most samples and maintaining sufficient sequencing depth for matched workflow comparison. Rarefaction remains debated in microbiome analysis because it can discard reads and reduce statistical efficiency, especially for differential abundance testing [19, 21]. This study used rarefaction only for like-for-like workflow comparison of alpha-diversity readouts, not for differential abundance testing or model-based taxon inference. This restricted use is consistent with reports that normalization choice depends on analysis goal and data characteristics [20]. Rarefaction can also remain defensible for community-level or ecological comparisons when interpreted with its known limitations [46]. These procedures were not intended to define a universal preprocessing recommendation.

### Nanopore analytical contexts

Two nanopore analytical contexts were considered for the paired n = 6 multi-platform dataset. The older nanopore taxonomic output was retained directly from the source-study analytical reference. The corresponding legacy platform environment had been discontinued and could not be re-executed. The newer nanopore output followed the current EPI2ME Desktop application (V5.3.1) using the latest ncbi_16s_18s reference database available in the default wf-16s analysis workflow.

Within-nanopore comparisons were therefore interpreted as comparisons between legacy reference-derived output and updated platform-native analysis. Cross-platform comparisons were not interpreted as pure instrument-only contrasts. They combined sequencing-platform and analysis-framework differences. The design could not decompose sequencing-platform effects from reference-database or algorithm-version effects. Nanopore old/new and cross-platform contrasts were therefore treated as compound analytical-context comparisons. These comparisons were retained to illustrate how platform, reference-database, and analysis-version differences can jointly affect diversity readouts, not to assign a separable effect size to any single technical factor. This whole-context comparison was used because the legacy nanopore environment could not be re-executed and a fully reference-matched platform-only contrast was not available.

### Zero handling and compositional transformation

Zero handling was performed before log-ratio transformation because microbiome count data are sparse and compositional. The main synbiotic and dietary-intervention Aitchison/CLR analyses used a pseudocount of 0.5 before CLR transformation. FeFiFo 16S rRNA gene amplicon analyses also used a pseudocount of 0.5, and FeFiFo CAZyme analyses used a pseudocount of 0.5 on processed CAZyme gene-family RPM abundance profiles. PRJEB41316 ASV/genus Aitchison outputs used a pseudocount of 1. The matched ASV/OTU workflow comparison relied on rarefied alpha-diversity and Bray-Curtis outputs rather than on a CLR/Aitchison workflow based on 97% OTU clustering. This pseudocount difference reflects analysis lineage and data-layer context rather than a biological threshold. The 0.5 pseudocount was used for the intervention response-geometry analyses. The pseudocount 1 PRJEB41316 output was limited to ASV/genus Aitchison support and not to the matched 97%-OTU coherence workflow.

The primary CLR representation was examined alongside an alternative Bayesian/multiplicative zero-replacement procedure, including cmultRepl-style replacement where available, as a sensitivity check [47, 48]. Where zCompositions::cmultRepl was unavailable, a custom deterministic Bayesian-multiplicative zero-replacement fallback was used only as a sensitivity implementation check, not as the primary representation. At a high level, the fallback assigned strictly positive mass to zero entries and rescaled nonzero entries to preserve compositional closure. Intervention response-geometry robustness runs also compared the primary 0.5 pseudocount with additional pseudocount settings where available, including half-minimum-positive and 1.0 settings.

CLR-transformed data were then used to represent each sample in compositional Euclidean space [49]. For paired analyses, each subject’s microbiome response was represented as a displacement from its own baseline sample to its corresponding follow-up sample within an Aitchison-based paired-response representation. The synbiotic and dietary-intervention worked example used the harmonized feature table from the primary analysis workflow without imposing an additional FeFiFo-style prevalence filter. The FeFiFo 16S rRNA gene amplicon analysis used a shared 5% prevalence filter across the requested timepoint-sensitivity sample set, as described below, to stabilize the FeFiFo feature space. This sample-specific baseline-referenced representation was used throughout the directional-coherence framework.

### Microbiome response magnitude

Microbiome response magnitude was defined as the size of paired compositional displacement between each subject’s baseline and post-intervention sample. Magnitude describes how far the microbiome moved from baseline. It was quantified in Aitchison-based compositional space using baseline-referenced paired distances, with larger values indicating greater overall microbiome displacement over the intervention interval. This quantity summarized response size rather than response orientation.

### Directional coherence analysis and diagnostic support

Directional coherence was used to assess whether paired microbiome responses within an intervention arm tended to align in a similar direction in an Aitchison-based ordination representation of compositional space [5, 50, 32]. Each subject contributed a baseline-referenced displacement vector defined from that subject’s own baseline sample to its follow-up sample within that representation [5, 50]. Coherence describes how similarly subjects moved in response space. It therefore reflects the organizational similarity of response directions across subjects, rather than the absolute location of samples in community space [4]. Cosine similarity was used as a scale-normalized directional summary, so coherence was evaluated together with response magnitude.

Magnitude and directional coherence were treated as complementary descriptors. A cohort or intervention arm could show modest average displacement but relatively coherent directionality, or larger displacement with weak directional agreement.

Permutation-based analyses evaluated whether observed directional structure exceeded random expectation [16]. These null procedures disrupted directional alignment while preserving the relevant paired-response structure. In sign-flip permutations, the unit was the participant-level paired response vector. Each vector was independently multiplied by +1 or −1 before recomputing the group-level coherence statistic; samples were not permuted across participants and group labels were not exchanged. This preserved vector length and the within-participant baseline-follow-up structure while disrupting consistent within-group orientation.

In the synbiotic and dietary-intervention worked example, the primary coherence statistic used the observed full-arm group mean direction. It was defined as the mean cosine similarity between each subject-level displacement vector and that group-level mean displacement vector. This calculation was performed in the Aitchison-based ordination representation. This convention follows the original Figure 4 implementation. Bootstrap resampling assessed the stability of group-level directional organization under modest perturbation. Leave-one-out influence checks were used as sensitivity analyses because group reference directions can be sensitive to individual response vectors in small groups. These checks are reported in Supplementary Table 5 as a dedicated leave-one-out influence sheet.

Because no single confirmatory primary endpoint or unified testing family was defined, p values were not pooled across the manuscript for a global multiplicity correction. Multiplicity adjustment was applied within local follow-up comparison sets where explicitly stated. Other p values were reported as analysis-specific permutation or robustness diagnostics. Exploratory pairwise arm comparisons for Figure 3 used Benjamini-Hochberg correction within that local comparison set. Directional-coherence permutation p values, sign-flip tests, implementation stress-test contrasts, and simulation detection summaries were reported as diagnostic or operating-behavior summaries rather than as one global confirmatory testing family.

**Figure 1.**
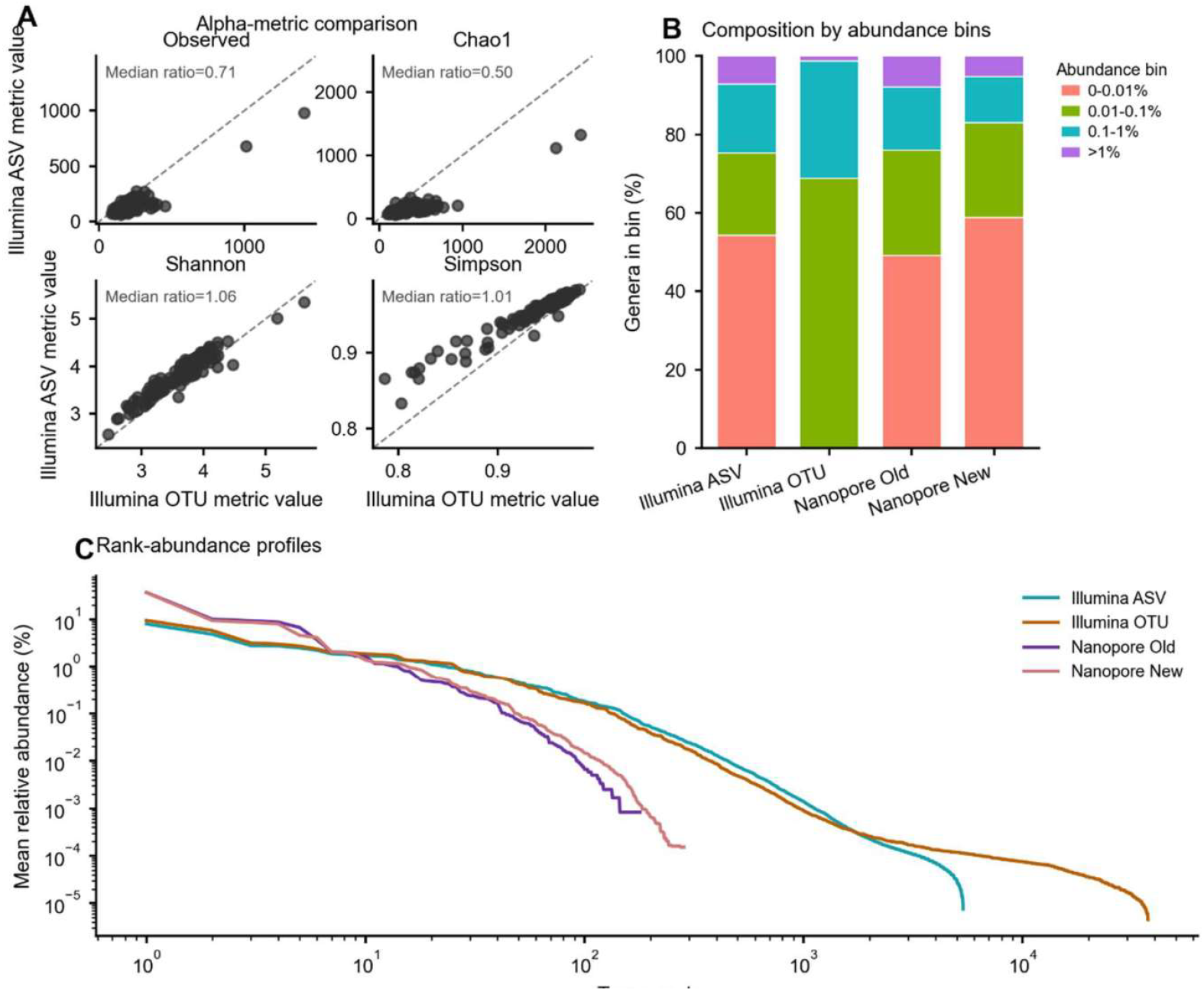
Richness and diversity inflation is concentrated in rare-tail structure and workflow-related abundance geometry. (A) Matched-sample Illumina OTU versus Illumina ASV alpha-diversity comparison for Observed richness, Chao1, Shannon, and Simpson; dashed diagonals indicate equality between workflows. Illumina OTU refers to the earlier OTU-based Illumina analysis workflow. (B) Composition by abundance bins across Illumina ASV, Illumina OTU, Nanopore Old, and Nanopore New using mean-abundance bins of 0-0.01%, 0.01-0.1%, 0.1-1%, and >1%. The absence of the 0-0.01% bin in Illumina OTU reflects workflow-specific clustering/filtering behavior rather than missing data. (C) Rank-abundance curves for the same workflow set on log-scaled rank and abundance axes.

**Figure 2.**
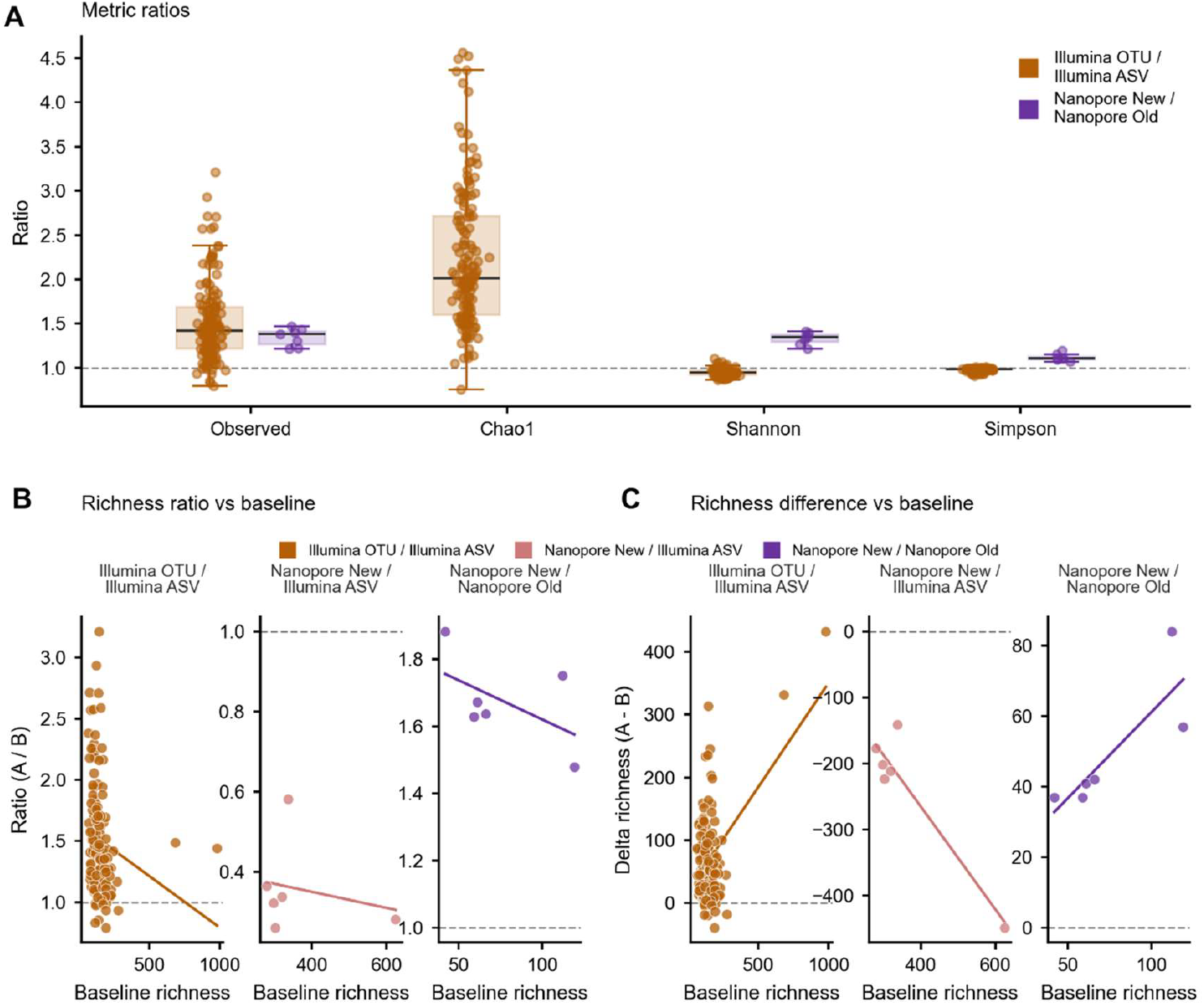
Workflow-associated richness inflation shows comparison-specific ratio and scaling behavior. (A) Metric-wise workflow ratios for the two within-platform preprocessing comparisons: Illumina OTU / Illumina ASV for Observed richness, Chao1, Shannon, and Simpson, and Nanopore New / Nanopore Old for Observed richness, Shannon, and Simpson only. Illumina OTU here denotes the earlier OTU-based Illumina analysis workflow. Nanopore Old denotes the legacy published nanopore genus table reported in the original study, whereas Nanopore New denotes the current EPI2ME rerun from public FASTQ files. (B) Multiplicative richness view showing the paired richness ratio A / B versus baseline richness for the three supported richness comparisons: Illumina OTU / Illumina ASV, Nanopore New / Illumina ASV, and Nanopore New / Nanopore Old. In the paired richness ratio A / B, A is the first workflow listed in the comparison title, B is the second workflow listed, and baseline richness denotes the richness value of workflow B. (C) Additive richness view showing the corresponding richness difference A - B versus baseline richness for the same comparisons. Here again, A is the first workflow listed in the comparison title, B is the second workflow listed, and baseline richness denotes the richness value of workflow B. Dashed horizontal lines indicate parity (A / B = 1) in panels A-B and zero difference (A - B = 0) in panel C.

**Figure 3.**
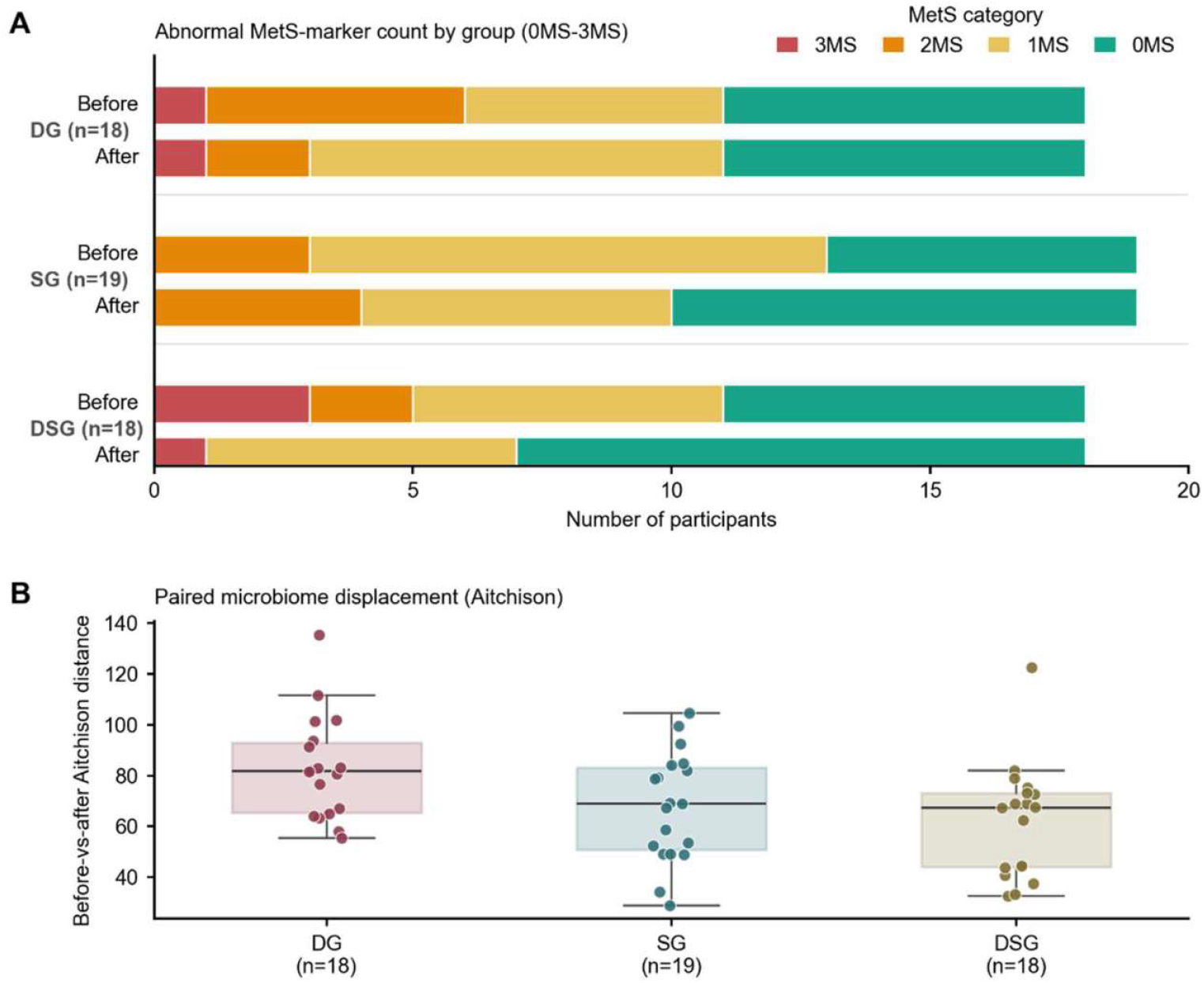
Clinical MetS burden and paired microbiome displacement across intervention groups. (A) Group-wise before/after distribution of abnormal MetS-marker count in paired participants from the dietary intervention group (DG, n = 18), synbiotic group (SG, n = 19), and combined diet-plus-synbiotic group (DSG, n = 18). 0MS, 1MS, 2MS, and 3MS denote the presence of 0, 1, 2, or 3 abnormal MetS-related markers among fasting glucose, HDL-C, and triglycerides; colors in Panel A encode these ordered categories. (B) Paired microbiome displacement magnitude shown as the before-versus-after Aitchison distance from the existing gold-trial paired microbiome distance table, derived from Euclidean distance in CLR-transformed microbiome space. Boxes show the interquartile range, center lines show medians, whiskers show the most extreme non-outlier values within 1.5 x IQR, and points represent individual participants. Inter-group differences in Panel B were assessed by Kruskal-Wallis test (p = 0.019); exploratory pairwise follow-up is summarized in the accompanying post hoc statistics file.

**Figure 4.**
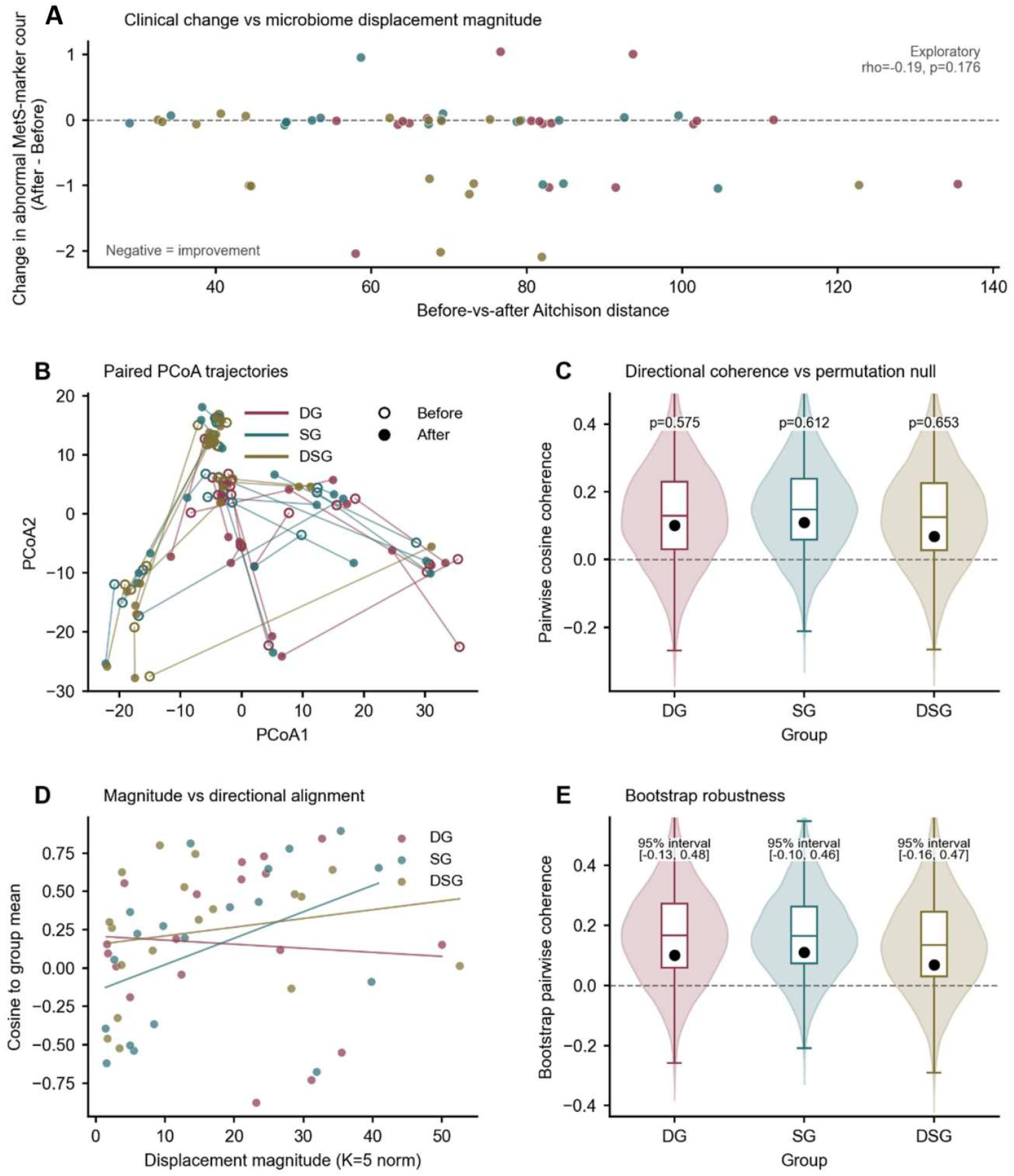
Direction-aware response analyses indicate weak within-arm directional coordination across intervention groups. (A) Exploratory association between paired microbiome displacement magnitude and change in abnormal MetS marker count (After - Before) in paired participants from DG (n = 18), SG (n = 19), and DSG (n = 18); negative values indicate improvement. Vertical jitter was applied to the y-axis for visualization, and the overall association was weak and not statistically supported (Spearman rho = −0.19, p = 0.176, n = 55). (B) Paired before/after ordination trajectories in Aitchison CLR-Euclidean space; open circles indicate before-intervention samples and filled circles indicate after-intervention samples. This panel is intended to visualize subject-level displacement heterogeneity rather than to imply a shared arm-level trajectory. (C) Group-level pairwise cosine coherence A compared with a within-subject before/after label-flip permutation null. Across arms, observed coherence remained close to the null reference, consistent with weak within-arm directional coordination in the current dataset. (D) Descriptive relation between displacement magnitude and alignment to the group mean direction. Lines are shown as descriptive visual guides only and should not be interpreted as formal slope inference. (E) Bootstrap distribution of the coherence statistic in CLR space. Confidence intervals broadly overlapped zero across arms, consistent with weak and uncertain within-arm directional coordination. Colors indicate intervention group across all panels.

The Figure 4 and Figure 5 directional-coherence summaries for the synbiotic and dietary-intervention cohort retained the full-arm mean direction to preserve the original Figure 4 implementation. Leave-one-out influence checks for this cohort are reported with the Figure 4 support in Supplementary Table 5 and were used to assess whether the full-arm convention was strongly driven by individual participants. The FeFiFo 16S rRNA gene amplicon, FeFiFo CAZyme gene-family, and stress-test analyses used leave-one-out mean directions to reduce self-influence in small groups. Both conventions address response-vector alignment, but absolute coherence values were not treated as directly interchangeable across datasets, endpoint definitions, or feature spaces.

**Figure 5.**
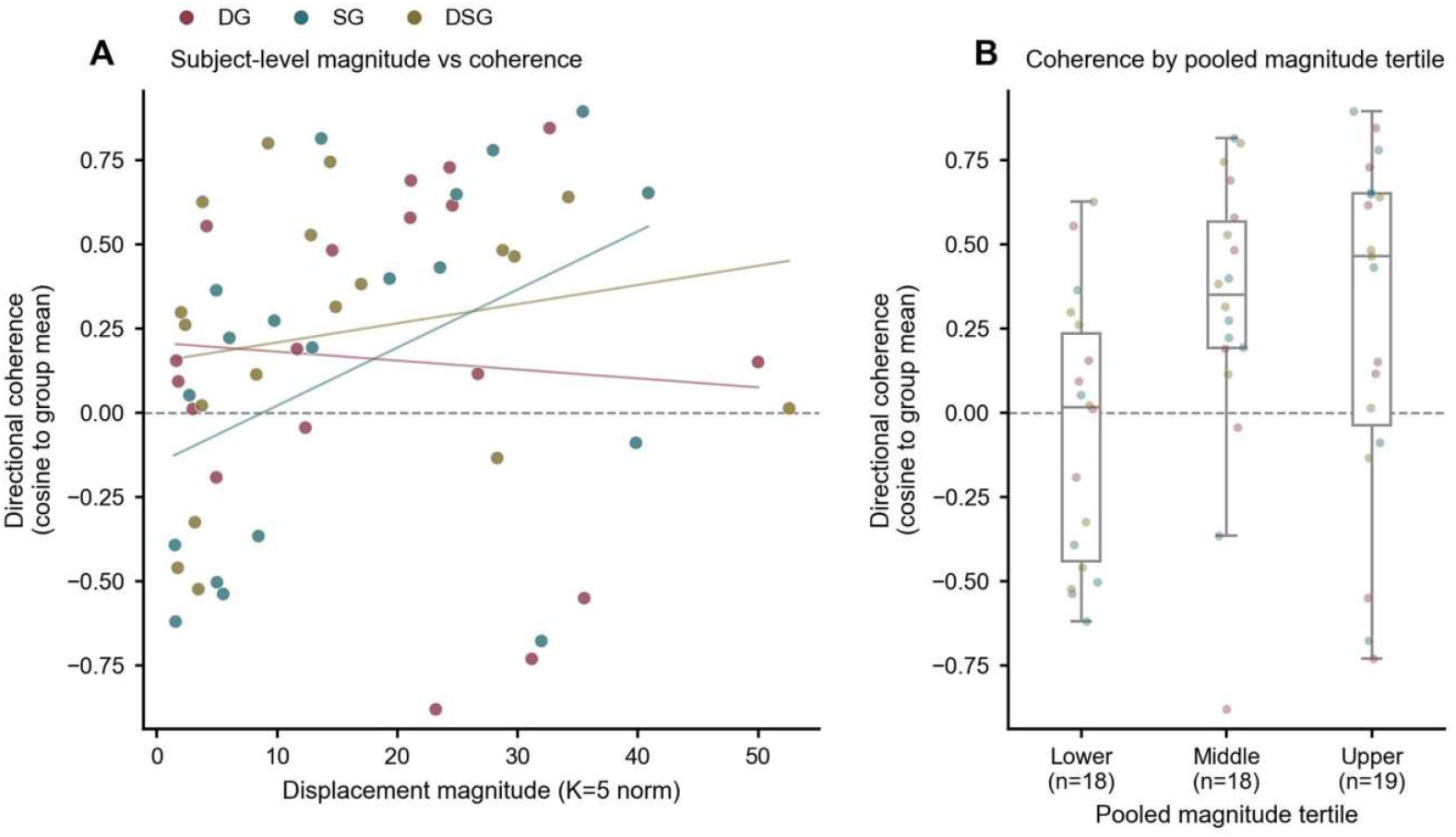
Displacement magnitude and directional coherence showed partial overlap but were not fully redundant in the intervention cohort. (A) Subject-level relation between K = 5 displacement magnitude and directional coherence, measured as cosine similarity to the corresponding group mean direction, for paired participants in DG (n = 18), SG (n = 19), and DSG (n = 18). Thin colored lines show within-group linear fits for visualization only. Across all subjects, the overall rank association was weak but positive (Spearman rho = 0.29, p = 0.033), whereas the group-adjusted linear slope was modest and not statistically supported (beta = 0.0069, p = 0.155; adjusted R2 = −0.01). (B) Coherence distributions across pooled magnitude tertiles, shown as a descriptive summary of spread at comparable displacement sizes. The pooled tertile view supports only partial overlap between response size and directional alignment rather than full redundancy.

### Rare-tail and scaling analyses

Richness changes across workflow transitions were examined with particular attention to low-abundance taxa because observed richness is sensitive to taxon detection, filtering, and sequencing-depth decisions [19, 15, 21]. Rare-tail analyses were used to determine whether richness increases were broadly distributed across the abundance spectrum or concentrated in the rare component of the community, where amplicon datasets may be more vulnerable to sequencing-introduced false positives and low-abundance feature instability [26, 27]. Shannon and Simpson results were used as abundance-weighted checks because these indices place more weight on abundance structure than observed richness alone [45, 15]. This approach helped separate rare-taxa expansion from broader changes in dominant community structure.

Paired comparisons were evaluated using both absolute differences and ratio-based behavior to characterize the geometry of workflow-induced diversity change. Workflow changes were summarized by asking whether diversity changed by a similar absolute amount across samples or scaled with the baseline diversity level.

Rare-tail operational sensitivity was further assessed using abundance-restriction and rare-tail attenuation analyses when suitable. These analyses tested whether workflow-associated richness contrasts weakened after lower-abundance taxa were removed or when analyses were restricted to dominant taxa. They were interpreted as operational sensitivity checks rather than as a universal abundance cutoff. This distinction was important because low-abundance taxa may contain ecological or host-relevant information [28], even though low-abundance amplicon features can also be more vulnerable to detection-boundary, sequencing-artifact, and filtering effects [26, 27].

### r500 simulation benchmark and implementation stress tests

An r500 simulation benchmark comprising 198,000 simulated datasets was used to examine the operating behavior of the coherence descriptor under null response, magnitude-only random-direction response, shared-direction response, mixed-responder response, opposing-responder subgroup, and rare-tail-sensitive scenarios. Synthetic baselines were generated from logistic-normal compositional profiles followed by multinomial sampling and zero handling; feature-dimensional settings included 100 and 500 features. Shared-direction effect-size settings were small = 0.35, medium = 0.8, and large = 1.4. The logistic-normal covariance structure was specified within the simulation grid rather than estimated from a specific empirical cohort, so the benchmark tested operating behavior under controlled compositional covariance settings rather than reproducing the covariance structure of any one dataset. Response scenarios imposed controlled response vectors in CLR space, consistent with the use of log-ratio geometry for compositional data [31, 22]. Shared-direction scenarios used increasing directional-effect settings, whereas magnitude-only random-direction scenarios increased displacement without imposing a common direction, reflecting the distinction between vector length and orientation in multivariate directional summaries [4, 5]. Full simulation parameter settings and values, including the baseline-generation model, zero-handling settings, feature counts, scenario definitions, effect-size grid, replicate structure, and summary outputs, are reported in Supplementary Table 8I.

The coherence gap was defined as observed group coherence minus the corresponding permutation-null mean. The benchmark was designed to test whether coherence remained near the null reference under random or magnitude-only response, increased under shared directional signal, weakened under mixed-responder behavior, and exposed cancellation under opposing-subgroup structure. Sample-size dependence was examined within the simulation grid rather than inferred from the small empirical cohorts alone. Rare-tail simulations were used to examine whether low-abundance feature inclusion could affect richness-sensitive summaries and whether dominant-abundance attenuation reduced apparent richness change.

Coherence gaps were interpreted relative to the sign-flip permutation null, bootstrap uncertainty, leave-one-out influence, and r500 simulation operating-reference summaries, including the nearest n = 20 simulation grid where relevant for the synbiotic and dietary-intervention arm sizes. These references were used as operating context rather than fixed universal thresholds, formal power calculations, or minimum detectable effects.

The simulation benchmark summarized expected analytical behavior and failure modes under controlled response scenarios. Empirical detection under null and magnitude-only scenarios was close to the nominal reference but slightly liberal in the aggregate grid, mainly in 100-feature settings. Detection in 500-feature simulations was closer to nominal. Full parameter grids, quality-control summaries, benchmark figures, and compact outputs are reported in Supplementary Table 8 and Supplementary Figures 6-9, with replicate-level outputs and provenance records retained in the GitHub reproducibility package.

Supplementary implementation stress tests were performed using David 2014 and Palleja 2018 public-data-derived analytical outputs [36, 37]. The David animal-diet run was used as a magnitude-stress contrast. Paired displacement magnitude, leave-one-out directional coherence, and sign-flip permutation support were calculated for the available Animal-arm anchor transition. This run assessed whether large displacement alone implied shared directional organization.

The Palleja antibiotic-recovery outputs were used as strong-perturbation implementation contrasts across species and pathway feature spaces. For each valid paired transition, subject-level response vectors were defined from the earlier timepoint to the later timepoint in CLR-transformed feature space. Response magnitude was calculated as the Euclidean length of each paired response vector. Leave-one-out directional coherence was calculated as cosine similarity to the same-transition group mean direction estimated without that subject. Sign-flip permutations preserving paired structure were used as diagnostic null support.

Disruption-versus-recovery cosines were calculated where endpoint structure allowed comparison of recovery vectors against early disruption vectors.

Pathway-space analyses used supplied pathway identifiers only. Magnitudes were interpreted only within feature space because species and pathway CLR coordinate systems are not scale-equivalent. These implementation stress tests are reported in Supplementary Table 9.

### Robustness, dispersion, non-redundancy, and exploratory clinical context

Additional robustness analyses were performed on existing data where appropriate. These included K-axis sensitivity, leave-one-out influence checks, high-magnitude or outlier exclusion, clinical-context robustness, rare-tail operational filtering sensitivity, and PERMANOVA/PERMDISP reference analyses. K-axis sensitivity refers to sensitivity to the number of retained PCoA/ordination axes used to represent Aitchison/CLR paired-response displacements before coherence calculation. These checks assessed whether the main response-organization interpretation was sensitive to representation choice, individual high-influence observations, or alternative multivariate summaries.

PERMANOVA and PERMDISP were used as complementary reference analyses when suitable. PERMANOVA provided a reference for centroid or group-location structure. PERMDISP-based analyses assessed whether apparent structure could be explained mainly by differences in multivariate spread rather than organized directional change [17]. These analyses were not treated as replacements for directional coherence. PERMANOVA and PERMDISP summarize centroid and dispersion structure. Directional coherence summarizes paired response-vector alignment.

The relationship between response magnitude and directional coherence was examined at the subject level. The analysis was summarized descriptively across magnitude strata to evaluate whether the two descriptors captured distinct aspects of microbiome change. This analysis summarized geometry and response organization.

Clinical and physiological variables in the synbiotic and dietary-intervention cohort, including abnormal MetS-marker transition context, were evaluated only as exploratory secondary analyses. These analyses examined whether microbiome response organization showed limited contextual correspondence with host outcomes in specific arms. FeFiFo host-context and functional layers were handled separately in the FeFiFo sections below.

Exploratory baseline-context analyses examined whether baseline microbiome features were associated with response magnitude or directional alignment within intervention arms. These analyses used the harmonized baseline feature table and were summarized as hypothesis-generating support in Supplementary Table 7D. They were not used for prediction, feature selection for the primary analyses, responder classification, or mechanistic inference.

### FeFiFo 16S rRNA gene amplicon application: endpoint design and preprocessing

The FeFiFo fiber/fermented-food dietary-intervention dataset provided an additional public intervention application of the magnitude-coherence framework, with high-fiber and high-fermented-food arms analyzed separately [2]. The 16S rRNA gene amplicon analysis used processed longitudinal phyloseq data rather than FASTQ reprocessing.

The primary 16S rRNA gene amplicon intervention endpoint was Baseline 0 to Week 14, corresponding to Timepoint 2 to Timepoint 9, because Week 14 was the final 16S rRNA gene amplicon endpoint available in the processed longitudinal object. This primary endpoint retained 17 fiber-arm and 16 fermented-arm pairs. Baseline −2 to Baseline 0, corresponding to Timepoint 1 to Timepoint 2, was used as a baseline-period negative-control interval and retained 18 pairs in each arm. Baseline −2 to Week 14, corresponding to Timepoint 1 to Timepoint 9, was retained as a maximum-span sensitivity interval and retained 17 fiber-arm and 16 fermented-arm pairs.

The FeFiFo 16S rRNA gene amplicon preprocessing workflow removed zero-count taxa, retained taxa present in at least 5% of the requested timepoint-sensitivity sample set, added a pseudocount of 0.5, and applied CLR transformation. In the FeFiFo 16S timepoint-sensitivity workflow, the prevalence sample set was defined as Fiber and Fermented 16S rRNA gene amplicon samples at the timepoints requested for the sensitivity contrasts. One shared feature set was used across requested pairs to keep trajectories comparable; the 5% filter was therefore not recalculated separately within each endpoint-specific paired sample set. The response vector for each participant and interval was defined as the final CLR profile minus the baseline CLR profile, and Aitchison magnitude was calculated as the Euclidean length of this displacement vector.

Directional coherence was summarized as leave-one-out cosine similarity to the group mean direction. For each participant, the reference direction was estimated from the remaining participants in the same arm to reduce self-influence. A sign-flip, paired-structure-preserving permutation was used as a diagnostic null. These analyses summarized response organization within each FeFiFo arm.

### FeFiFo CAZyme gene-family response-space coherence

A constrained CAZyme gene-family response-space analysis examined whether the framework provided functional-context information in a metagenomics-derived gene-abundance feature space. CAZyme gene-family abundance profiles were derived from shotgun metagenomic analysis of the FeFiFo study and were not inferred from 16S rRNA gene amplicon data. These profiles represent metagenomic CAZyme gene-family abundances, not measured enzyme activity.

The CAZyme analysis used 840 retained metagenomics-derived glycoside hydrolase and polysaccharide lyase (GH/PL) CAZyme gene-family RPM feature IDs, comprising 793 GH and 47 PL features represented at family-plus-subfamily level. This retained gene-family feature space was not restricted to the 11 CAZyme genes reported as significantly increased in the original high-fiber analysis. The CAZyme gene-family response vector for each participant was defined as the endpoint profile minus the corresponding baseline profile after CLR transformation. Functional-context organization was operationalized as leave-one-out directional coherence in this retained CAZyme gene-family response space. A sign-flip, paired-structure-preserving permutation was used as a diagnostic null. The leave-one-out mean direction was estimated within each arm.

A targeted CAZyme gene-family feature-retention sensitivity check compared the retained GH/PL gene-family feature space with alternative feature-retention rules. The targeted checks included a stricter prevalence rule and reduced feature-space subsets based on abundance or variance, as summarized in Supplementary Table 13E. These checks assessed feature-space dependence rather than providing an exhaustive search for an optimal CAZyme filtering rule.

The CAZyme gene-family endpoint followed the original-paper-compatible baseline-to-end-maintenance contrast because the processed CAZyme gene-family profiles did not include the exact Week 14 endpoint used for the primary 16S rRNA gene amplicon analysis. Available CAZyme profiles used Timepoint 1 / Baseline - 2 as the baseline when available, otherwise Timepoint 2 / Baseline 0. The endpoint was Timepoint 7 / Week 10 when available, otherwise Timepoint 6 / Week 8. This timing differs from the primary 16S rRNA gene amplicon Week 14 endpoint. The comparison was retained because it used measured metagenomics-derived CAZyme gene-family abundance profiles from the same dietary-intervention study and provided a constrained functional-context view of response organization. The non-identical timing was recorded as an endpoint caveat for cross-layer interpretation. Because no exact matched 16S rRNA gene amplicon and CAZyme gene-family endpoint pair was available in the processed objects used here, no matched-endpoint cross-layer inference was attempted. CAZyme gene-family preprocessing, endpoint mapping, and directional-coherence calculations are reported in Supplementary Tables 12 and 13.

### Software environment and supplementary framework extension

Analyses were conducted primarily in R (version 4.4.2) using the packages dada2 (1.34.0), phyloseq (1.41.1), and vegan (2.7-2). VSEARCH [44] was used for OTU-oriented clustering. EPI2ME Desktop (V5.3.1) was used for the updated nanopore workflow. Additional R and Python scripts were used for figure generation, r500 simulation benchmarking, David/Palleja implementation stress tests, robustness summaries, and supplementary table assembly. Python version and package details are reported in the accompanying code logs where applicable.

Reporting and reproducibility practices were aligned with contemporary microbiome-study recommendations emphasizing accurate terminology, detailed wet-lab and in silico workflow description, and data, protocol, and code availability [51].

A supplementary script-based implementation of the Figure 7 magnitude-coherence framework is provided as a beta research-preview GitHub repository at https://github.com/carolyyszeto/microbiome-response-interpreter-beta, tagged as v6.5-beta. The repository supports exploratory replication, toy-data demonstration, output interpretation, pseudocount sensitivity checks, simulation-based operating-context exploration, and descriptive application. This implementation is intended as a manuscript-aligned companion workflow rather than a validated clinical, diagnostic, predictive, efficacy, regulatory, production, or mechanistic software tool.

**Figure 6.**
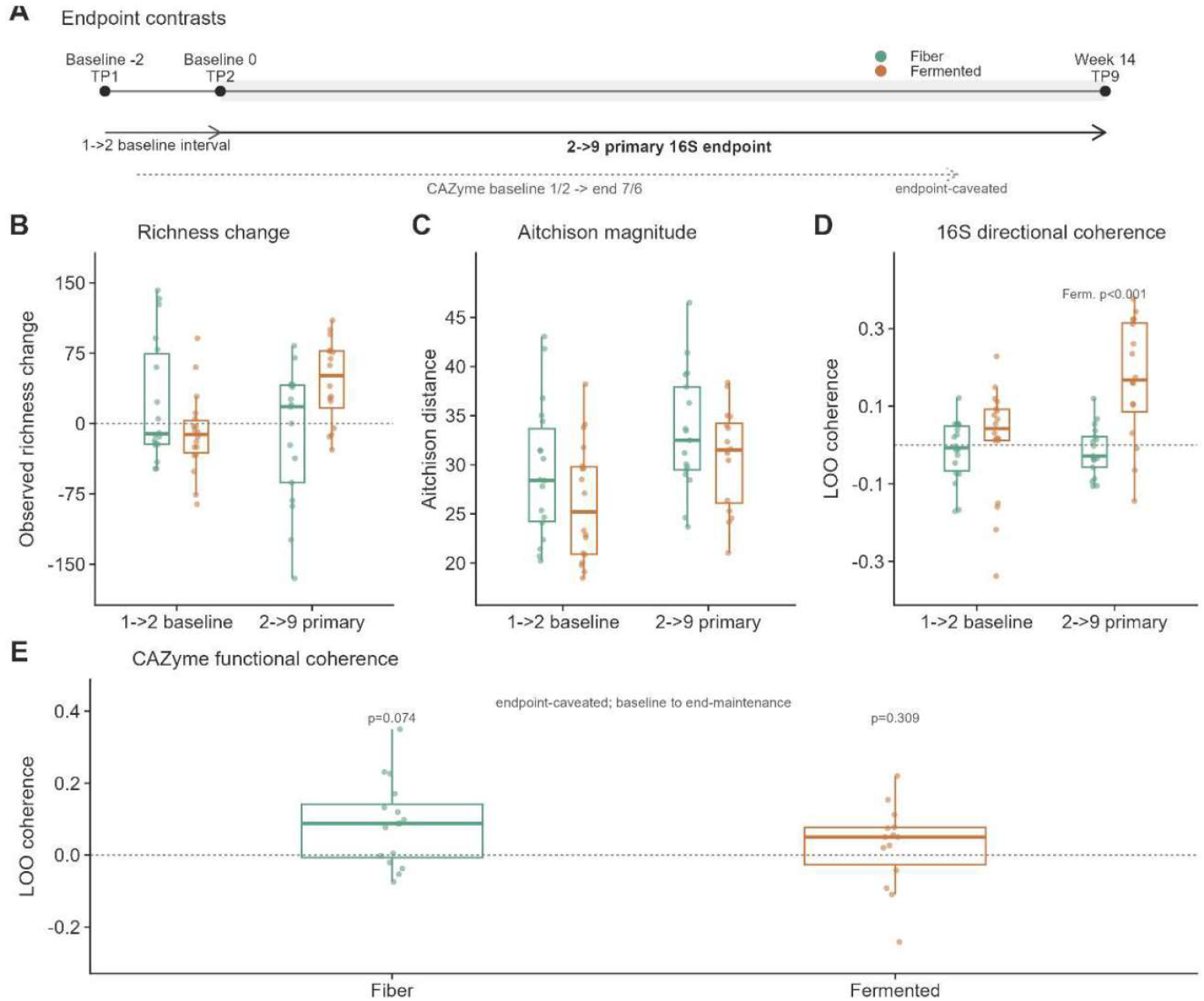
FeFiFo fiber/fermented-food dietary-intervention separates taxonomic and functional response organization. (A) Endpoint contrasts used for the Wastyk fiber/fermented-food application. The 1->2 contrast represents the Baseline −2 to Baseline 0 interval, the 2->9 contrast represents the primary 16S rRNA gene amplicon endpoint from Baseline 0 to Week 14, and the CAZyme contrast represents the available baseline-to-end-maintenance functional endpoint. (B) Subject-level observed-richness change across the 1->2 and 2->9 16S contrasts in the Fiber and Fermented arms. Points denote paired participants, boxes show interquartile ranges, center lines show medians, and the dashed horizontal line indicates no change. (C) Subject-level Aitchison displacement magnitude across the same 16S contrasts. This panel summarizes the size of microbiome movement in compositional space. (D) Subject-level leave-one-out directional coherence across the same 16S contrasts. Directional coherence was measured as the cosine similarity between each subject’s response vector and the group-mean response direction estimated without that subject. Sign-flip permutation p values shown in the panel summarize group-level support for directional coherence and are not subject-level p values. (E) CAZyme functional feature-space directional coherence in the Fiber and Fermented arms using the baseline-to-end-maintenance contrast. Points denote paired participants, boxes show interquartile ranges, center lines show medians, and the dashed horizontal line indicates zero coherence. The CAZyme endpoint is not identical to the 16S Week 14 endpoint and is therefore interpreted as endpoint-caveated functional context.

**Figure 7.**
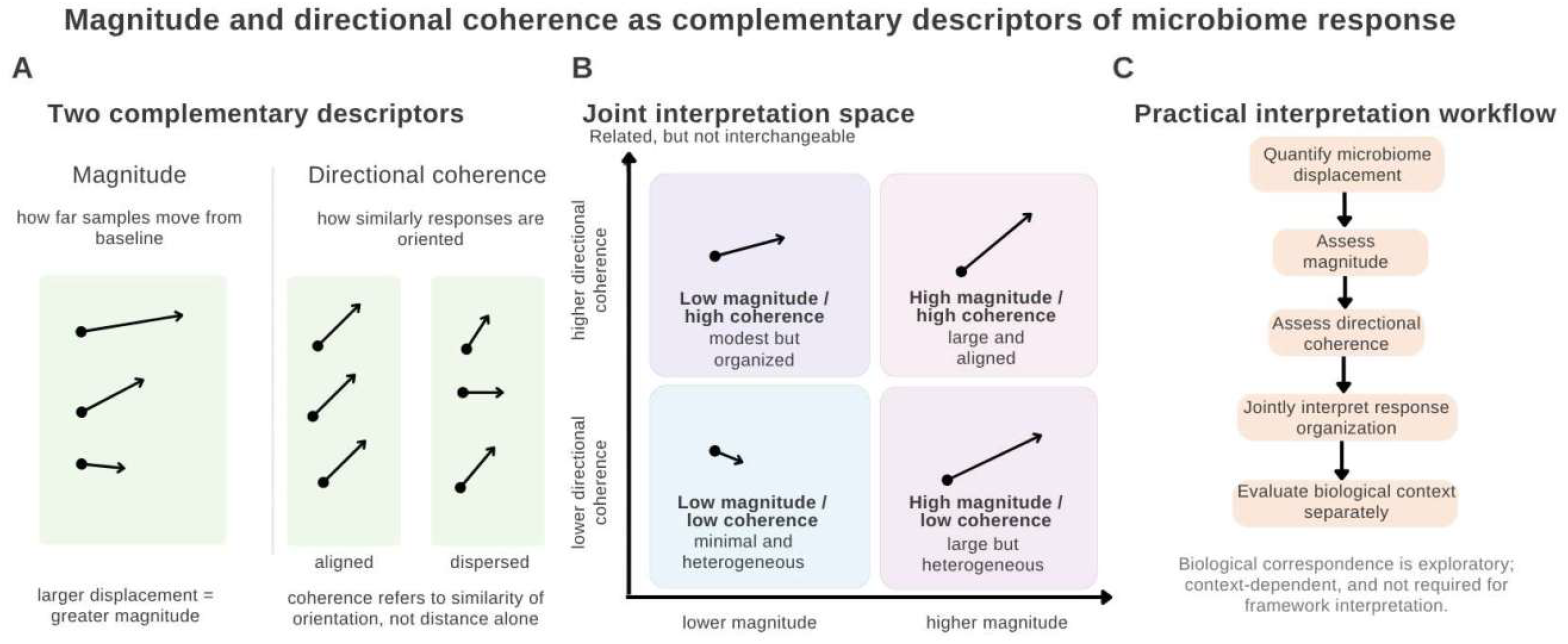
Conceptual framework for interpreting microbiome response across analytical context, response magnitude, and directional coordination. (A) Response magnitude is defined as displacement from each sample’s baseline reference and directional coherence as similarity in response orientation. (B) A conceptual joint interpretation space is shown with four response states across low- and high-magnitude settings. The boundaries are conceptual rather than empirical and are intended to guide interpretation rather than assign samples to discrete categories. In the present intervention setting, the overall pattern was more consistent with heterogeneous response organization than with a strongly aligned response state. (C) A general interpretation workflow is outlined in which microbiome displacement is quantified, magnitude and directional coordination are assessed, response organization is interpreted jointly, and downstream biological context is considered separately. Biological or physiological correspondence remains secondary and exploratory rather than being required for framework interpretation.

### Use of large language model assistance

Several large language model tools were used to assist with manuscript drafting and language editing. These tools were not used as authors and were not used to generate, analyze, or independently verify the primary data. All analyses, code outputs, figures, supplementary tables, citations, interpretations, and final manuscript text were reviewed and approved by the authors. The authors take full responsibility for the integrity and accuracy of the work.

## Results

The Results proceed from analytical sensitivity to operating behavior and then to empirical intervention applications. Figures 1 and 2 examine workflow-sensitive richness behavior. The r500 simulations and David/Palleja stress tests define operating behavior and empirical operating contrasts. Figures 3-5 apply the framework to the synbiotic and dietary-intervention cohort, and Figure 6 applies it to the FeFiFo dietary-intervention dataset.

### Workflow sensitivity concentrated in rare-tail richness structure

Figures 1 and 2 first examined how analytical workflow affected richness and diversity readouts in the Santiago-Olesen Illumina stool-donor cohort and the Matsuo paired nanopore/Illumina dataset [39, 40]. This comparison was framed within the broader context that microbiome metric choice and analytical workflow can materially affect community interpretation [18]. The main workflow effect was concentrated in richness-oriented summaries and low-abundance community structure, rather than affecting all diversity measures equally.

In the matched Illumina OTU-versus-ASV comparison, Observed richness and Chao1 separated more strongly than Shannon or Simpson (Figure 1A). This pattern was consistent with greater sensitivity of richness-oriented metrics to low-abundance features. The broader workflow comparison localized the largest qualitative differences to the lowest-abundance bin (0-0.01%) (Figure 1B). This bin contained 114 genera in Illumina ASV and none in Illumina OTU, compared with 88 in Nanopore Old and 167 in Nanopore New. The missing Illumina OTU bin reflects workflow-specific clustering or filtering rather than missing data. Nanopore New also retained more very low-abundance taxa than Nanopore Old. Patterns outside the lowest bin were heterogeneous rather than uniformly attenuated (Supplementary Table 1).

Rank-abundance structure provided a complementary view of the same workflow sensitivity (Figure 1C). In the shared 0.1-1% mid-to-low abundance bin, Illumina ASV showed the highest mean richness proportion, Illumina OTU showed the lowest value, and the nanopore workflows were intermediate (Supplementary Table 1). Attenuation analyses further supported rare-tail sensitivity in the Illumina ASV-to-OTU comparison. The full observed delta mean was 210.318 in the full matrix, but it reversed sign to −0.25 after restriction to the top 10 taxa. The delta mean also remained negative at −9.574 after restriction to taxa accounting for the top 80% of cumulative abundance. Together, these results were consistent with rare-tail sensitivity in the full-matrix richness contrast rather than a uniform change across dominant community structure (Supplementary Table 3).

Figure 2 showed that workflow-associated richness differences also followed comparison-specific ratio and scaling patterns. In Figure 2A, the largest departures from parity occurred in richness-oriented summaries. The Illumina OTU-versus-ASV median ratios were 1.418 for Observed richness and 2.01 for Chao1, compared with 0.945 for Shannon and 0.986 for Simpson. Chao1 was retained in the Illumina OTU-versus-ASV panel as a singleton-sensitive richness stress case, not as a cross-platform endpoint. This distinction is important because Chao1 is a singleton-dependent richness estimator that is difficult to compare across workflows with different rare-feature handling [52]. Chao1 was not included in the legacy Nanopore Old pipeline. The Nanopore New-versus-Old comparison was therefore limited to the matched set of Observed richness, Shannon, and Simpson. Within this matched set, the median ratios were 1.379 for Observed richness, 1.348 for Shannon, and 1.108 for Simpson.

Figure 2B showed that richness ratios did not follow a single common scaling pattern across comparisons. The Illumina OTU/ASV ratio was higher at lower baseline richness and moved toward parity as baseline richness increased, although the overall rank correlation remained near zero (rho = −0.034, p = 0.68). Nanopore New/Old ratios were concentrated over a narrower baseline-richness range and remained mostly above parity. The Nanopore New-versus-Illumina ASV comparison had only six matched samples. It remained below parity across the observed range and was treated as exploratory scaling context only (rho = −0.371, p = 0.468; Supplementary Table 2).

Figure 2C showed that additive richness differences were also comparison-specific. The fitted lines were used as simple scaling summaries, not as models intended to capture all local pattern changes. This distinction was most important for the Illumina OTU-versus-ASV contrast, where the fitted slope was positive but the plotted points showed a sign reversal. OTU-ASV differences were positive at lower baseline richness and negative at higher baseline richness. The Nanopore New-versus-Old contrast also showed a positive scaling direction. The small cross-platform Nanopore New-versus-Illumina ASV contrast showed a negative scaling direction, but this n = 6 comparison was used only as exploratory scaling context. All model intercepts were extrapolative because observed baseline richness did not extend to zero. The corresponding estimates are reported in Supplementary Table 2.

Rare-tail-sensitive richness shifts were evaluated using attenuation rather than a universal abundance cutoff. Here, a rare-tail-sensitive classification means that the observed-richness contrast markedly weakened after dominant-only or cumulative-abundance restriction. Supplementary Table 3 contains attenuation support for both the Illumina ASV-to-OTU comparison and the Nanopore New-versus-Old comparison. In the nanopore support analysis, the full-matrix richness difference was 111.6, the top-20 difference was 1.43, and the top-20 attenuation value was 0.987. This supported interpreting the nanopore comparison as rare-tail-sensitive within the present attenuation framework, while the operational classification was not generalized beyond the tested comparisons.

### Simulation benchmarking defined operating behavior

The r500 simulation benchmark evaluated the operating behavior of the response-geometry framework before interpretation of empirical intervention datasets. The benchmark included 198,000 simulated datasets spanning null-response, magnitude-only random-direction, shared-direction, mixed-responder, opposing-subgroup, and rare-tail-sensitive scenarios (Supplementary Table 8; Supplementary Figures 6-9).

Null-response simulations showed coherence gaps centered near zero. Magnitude-only random-direction simulations also remained near the null reference. These two negative-control settings showed that random movement or larger displacement alone was not interpreted as coordinated directional organization (Supplementary Table 8; Supplementary Figures 6 and 8).

Shared-direction simulations provided the corresponding positive-control behavior. Coherence gaps and detection rates increased with stronger shared-direction effects and larger sample sizes. This pattern indicated that the coherence descriptor increased when a shared response direction was present. Mixed-responder simulations reduced pooled coherence because only a subset of participants followed the shared direction. Opposing-subgroup simulations showed that high displacement could coexist with low pooled coherence when response directions cancelled across subgroups (Supplementary Table 8; Supplementary Figures 6-8).

Rare-tail simulations addressed the richness-sensitivity component of the framework. Richness-oriented summaries were strongly affected by inclusion of low-abundance features, whereas dominant-abundance attenuation markedly reduced apparent richness change. This result was consistent with the rare-tail attenuation logic used in Figures 1 and 2, while keeping the simulation benchmark separate from biological interpretation (Supplementary Table 8; Supplementary Figure 9). The slight liberal behavior observed in the benchmark was concentrated mainly in the 100-feature setting. Accordingly, p values from smaller filtered feature spaces should be interpreted as diagnostic operating-context information rather than stand-alone confirmatory evidence.

### Strong-perturbation implementation stress tests provided operating contrasts

David/Palleja implementation stress tests provided empirical operating contrasts under stronger perturbation and magnitude-stress settings [36, 37]. The dataset roles and claim boundaries are summarized in Supplementary Table 9C. These analyses were used as implementation stress tests only. They were not interpreted as biological validation, clinical evidence, participant classification, or general framework performance.

The Palleja antibiotic-recovery trial-run outputs served as a qualitative strong-perturbation implementation contrast. Early disruption transitions showed positive directional organization in both species and pathway feature spaces (Supplementary Table 9A). Recovery vectors were reversal-like relative to early disruption vectors, a pattern consistent with endpoint-aligned perturbation followed by recovery-oriented movement (Supplementary Table 9B). This contrast does not define an effect-size operating range for dietary-intervention applications.

The David animal-diet trial-run output served as the magnitude-stress contrast. The available Animal-arm anchor transition showed large displacement, but it did not show clear leave-one-out directional organization above the sign-flip null (Supplementary Table 9A). This contrast separated large movement from shared directionality and was consistent with the interpretation that displacement magnitude alone is insufficient for inferring coordinated response geometry.

### Intervention magnitude distributions were modest and heterogeneous across arms

Figure 3 applied the framework to the synbiotic and dietary-intervention cohort [38]. This cohort was used as a modest, heterogeneous intervention example rather than as clinical-use evidence. The figure summarized abnormal MetS-marker count transitions and paired microbiome displacement across the dietary intervention group (DG; n = 18), synbiotic group (SG; n = 19), and combined diet-plus-synbiotic group (DSG; n = 18).

Figure 3A showed the clearest post-intervention redistribution toward lower abnormal MetS-marker count categories in DSG. Participant-level transition support showed that 17 of 18 DSG participants were in the 0MS or 1MS categories after intervention, compared with 15 of 19 in SG and 15 of 18 in DG. DSG also showed the largest number of downward category shifts, with 8 of 18 participants shifting downward, compared with 4 of 18 in DG and 3 of 19 in SG (Supplementary Figure 1; Supplementary Table 4). Figure 3A should therefore be read as exploratory clinical-context information, not as evidence of uniform response within every arm.

Figure 3B showed that paired microbiome response magnitude differed across groups. The overall Kruskal-Wallis test was statistically significant (p = 0.019). DG and DSG were separated after Benjamini-Hochberg correction in exploratory pairwise follow-up (raw p = 0.008; adjusted p = 0.023). DG versus SG and SG versus DSG were not clearly separated after correction, leaving SG intermediate in this magnitude comparison (Supplementary Table 4). The arm with larger global taxonomic displacement was therefore not the arm with the clearest descriptive MetS-marker redistribution.

### Directional-coherence analyses showed weak within-arm coordination

Figure 4 asked whether intervention-associated microbiome shifts were directionally organized within arms after response magnitude was quantified. The primary Figure 4 analysis used the K = 5 Aitchison CLR-Euclidean representation, with K-sensitivity summarized in Supplementary Figure 3 and Supplementary Table 5. The main finding was weak within-arm directional coordination in a cohort with substantial subject-level displacement heterogeneity.

Figure 4A provided an exploratory link between paired microbiome displacement magnitude and change in abnormal MetS-marker count. The association was weak and not statistically supported (Spearman rho = −0.19, p = 0.176, n = 55). This result suggested that displacement magnitude alone did not track clinical improvement in a simple monotonic manner at the cohort level.

Figure 4B visualized paired before-after paths in Aitchison CLR-Euclidean space. These paths were heterogeneous across participants and did not suggest a shared arm-level directional pattern by visual inspection. Figure 4C then evaluated this visual impression using group-level directional coherence. In Figures 4 and 5, group-level directional coherence refers to cosine alignment between each subject-level response vector and the corresponding group mean direction. Higher values indicate that participants moved more similarly in compositional response space. Observed group-level directional coherence was compared with a within-subject before-after label-flip permutation null. This null preserved paired structure while disrupting consistent directional alignment.

Observed group-level directional coherence remained close to the permutation-null reference across DG, SG, and DSG. The corresponding permutation p values were 0.575, 0.612, and 0.653, respectively (Supplementary Table 5). These results did not provide evidence for a strong shared microbiome response direction within any intervention arm. The current sample size also limited detection of moderate shared-direction effects if present.

Figure 4D provided a descriptive check of whether larger displacement implied stronger directional alignment at the subject level. Figure 4E addressed precision by bootstrap resampling. The CLR-space bootstrap distributions broadly spanned zero across arms. K-sensitivity, permutation sensitivity, bootstrap robustness, and secondary divergence support did not change the main interpretation of weak within-arm coordination (Supplementary Figures 2-5; Supplementary Tables 5 and 6).

Zero handling is part of the CLR response-vector definition. A targeted pseudocount sensitivity check was performed under fixed sample pairing, endpoint definitions, and feature filters. In the Figure 4 analysis, rerunning the directional-coherence calculation with a pseudocount of 1.0 did not change the qualitative calls for DG, SG, or DSG compared with the primary 0.5 setting. These results reduced concern that the Figure 4 interpretation was driven by the single 0.5 pseudocount choice, although they do not establish invariance to pseudocount choice or replace a full zero-handling benchmark (Supplementary Table 5).

Clinical-context robustness analyses were consistent with the same interpretation in this modest sample. The association between displacement magnitude and abnormal MetS-marker change remained weak, and bootstrap and permutation checks did not provide stronger support for a monotonic clinical-context relation. Arm-specific estimates were also imprecise, with wide intervals that crossed zero. SG had more participants with low MetS-marker burden at baseline, leaving less room for MetS-marker improvement than in the other arms. This baseline imbalance limited clinical-context interpretation rather than establishing a stronger or weaker microbiome directional response. Numerical robustness summaries are provided in Supplementary Table 5.

### Response magnitude and directional coherence were partially overlapping but not redundant

Figure 5 examined whether response magnitude and directional coherence behaved as redundant descriptors of the same response property. Magnitude and directional alignment showed partial overlap, but the relation was weak.

In Figure 5A, subjects with larger displacement tended to rank with higher directional alignment across the pooled cohort (Spearman rho = 0.29, p = 0.033; Supplementary Table 7A). The association was not supported as a simple group-adjusted linear trend (beta = 0.0069, 95% CI −0.0027 to 0.0165, p = 0.155; Supplementary Table 7B). The group-adjusted model added little explanatory value (adjusted R2 = −0.008), and an interaction test did not support different magnitude-coherence slopes across groups (p = 0.247). The current sample size limited detection of small or subgroup-specific relationships. These analyses were consistent with partial overlap between response size and directional alignment, but not with a strong shared response descriptor.

Figure 5B illustrated why the two descriptors were not redundant. Directional alignment increased from the lower to the upper pooled magnitude tertiles, but each tertile still showed broad spread. Subjects with similar displacement size could therefore differ substantially in directional alignment. No formal trend test was applied because the tertile view was descriptive and each bin retained mixed group membership. The corresponding tertile summary is provided in Supplementary Table 7C.

Exploratory baseline-context analyses added arm-specific biological context without modifying the primary Figure 5 analysis. Baseline microbiome features were most consistently associated with the magnitude-coherence descriptors in SG, whereas DG and DSG showed largely weak or null baseline patterns. This arm-level contrast suggested that baseline microbiome state may have been more relevant to response organization in SG than in the other arms. These observations were not used to modify the primary analysis strategy and were not treated as participant-classification or causal results (Supplementary Table 7D).

### FeFiFo dietary-intervention analysis showed distinct taxonomic and timing-limited functional response patterns

The FeFiFo dataset was used as a second dietary-intervention application after the synbiotic and dietary-intervention cohort [2]. This analysis applied the same response-organization descriptors separately to 16S rRNA gene amplicon taxonomic response space and to endpoint-caveated CAZyme gene-family feature space. It did not test matched taxonomic-functional coupling or re-test the original FeFiFo primary outcomes.

Figures 6A-D summarized 16S rRNA gene amplicon response organization. The Baseline −2 to Baseline 0 interval (1->2) served as a baseline-period control, and the Baseline 0 to Week 14 interval (2->9) served as the primary 16S rRNA gene amplicon endpoint (Figure 6A; Supplementary Tables 10 and 11; Supplementary Figure 10). In the baseline-period control, leave-one-out coherence was near zero in both arms: −0.015 in Fiber (permutation p = 0.712) and 0.012 in Fermented (permutation p = 0.343). These baseline-period values were close to the sign-flip null reference. Fermented-food exposure showed a stronger richness increase and positive leave-one-out directional coherence at the primary endpoint (Figures 6B and 6D). High-fiber exposure did not show comparable 16S rRNA gene amplicon taxonomic directional coherence (Figure 6D). Figure 6C shows displacement magnitude, whereas Figure 6D shows directional organization. These panels therefore separate movement size from response organization in the same taxonomic feature space.

Figure 6E applied leave-one-out directional coherence to metagenomics-derived CAZyme gene-family profiles. In this analysis, each participant was compared with the group mean direction estimated from the other participants, reducing self-influence in small arms. The analysis used 840 retained CAZyme RPM feature IDs in the GH/PL feature space rather than only the 11 CAZyme features reported as increased in the original high-fiber analysis. This 840-feature CAZyme space was defined as the retained metagenomics-derived GH/PL RPM feature set available for paired response-vector analysis, rather than being selected from the 11 originally increased CAZymes. The retained CAZyme feature space therefore evaluated multivariate CAZyme profile organization rather than specific abundance-change features from the original report. This distinction matters because the original upregulation analysis asked which CAZyme features increased in abundance, whereas directional coherence asked whether participant-level CAZyme profiles moved in a similar multivariate direction within the retained feature space.

The CAZyme analysis used the available baseline-to-end-maintenance contrast. Exact time points varied with metagenomic sample availability, using Baseline −2 or Baseline 0 as baseline and Week 10 or Week 8 as endpoint. This timing was not identical to the primary 16S rRNA gene amplicon endpoint from Baseline 0 to Week 14. The timing mismatch may affect coherence estimates and prevents a direct matched-endpoint comparison at the primary 16S Week 14 endpoint. The CAZyme result was therefore retained as endpoint-caveated functional-context support, not as primary evidence for a matched cross-layer conclusion.

Within the original timing-mismatched contrast, the fiber arm showed weak-to-modest CAZyme-gene directional coherence, whereas the fermented arm remained closer to zero. Arm-level magnitudes, coherence estimates, paired sample sizes, and sign-flip permutation results are provided in Supplementary Figure 11 and Supplementary Tables 12 and 13. These results were retained as descriptive response-organization findings. The non-identical CAZyme endpoint prevents direct cross-layer comparison, and the CAZyme-gene result should be interpreted within its measured feature space. It should not be interpreted as a measure of catalytic function, metabolic flux, or mechanism.

An endpoint-aligned sensitivity check tested endpoint dependence near the Figure 6E interpretation using the best-supported CAZyme-aligned endpoint, Baseline 0 to Week 8 (T2 to T6). This check retained CAZyme paired samples for Fiber n = 12 and Fermented n = 11, and an exact-subject 16S rRNA gene amplicon rerun was performed on the same participants. The Baseline 0 to Week 8 matched-timepoint and exact-subject rerun did not reproduce the original Fiber-dominant CAZyme coherence pattern. CAZyme mean leave-one-out coherence was 0.004 in Fiber and 0.092 in Fermented, while exact-subject 16S rRNA gene amplicon values were 0.020 and 0.085, respectively. CAZyme coherence was therefore interpreted as endpoint-sensitive, feature-space-dependent functional-context support (Supplementary Table 13C).

Targeted pseudocount sensitivity checks also supported the conservative FeFiFo interpretation. The 0.5 versus 1.0 comparison did not change the qualitative interpretation of either the 16S primary endpoint or the endpoint-caveated CAZyme analysis. The CAZyme Fiber estimate remained modest and borderline under both settings. In the optional endpoint-matched CAZyme check, the Fiber estimate shifted from a tiny positive value to a slightly negative value, reinforcing that the aligned-endpoint CAZyme result should be treated as sensitivity-dependent contextual support rather than stable functional response organization (Supplementary Table 13).

## Discussion

Dietary and fiber-focused microbiome interventions often produce mild or layer-specific remodeling rather than uniform large community shifts [1, 2]. Probiotic responses can also vary across individuals [6]. These responses can be heterogeneous and baseline-dependent [1, 6, 8]. Scalar diversity or group-level summaries can therefore appear weak or inconclusive. This can occur when participants show organized but magnitude-limited movement, or when they move substantially in divergent directions [4, 5]. Microbiome diversity remains useful for summarizing community change, but its interpretation can depend strongly on analytical context. Alpha-diversity and beta-diversity summaries describe related but distinct aspects of community structure, including within-sample richness or evenness, between-sample dissimilarity, and multivariate dispersion [10, 11, 3]. Workflow differences can reshape richness-based readouts without equivalent effects on abundance-weighted summaries, complicating comparisons across datasets and processing backgrounds. The same issue extends to beta-space interpretation, because metric choice, transformation, and feature-space definition shape how microbiome movement is represented [18]. These sensitivities create a need to separate what analytical choices change from what biological perturbation moves within a defined analysis space.

This study addresses that need through a response-geometry framework for microbiome remodeling during intervention periods. Most applications in this study involved limited and heterogeneous responses rather than uniform large perturbations. Response magnitude describes how far each microbiome profile moved from baseline. Directional coherence describes whether participant-level response vectors were oriented in similar directions. This distinction is consistent with trajectory-based community ecology, in which path length and trajectory direction can represent different dimensions of temporal community change [5]. These layers therefore distinguish workflow-driven diversity shifts from microbiome displacement and separate heterogeneous movement from coordinated participant-level response. Figure 7 summarizes this logic by positioning diversity sensitivity, response magnitude, directional organization, and downstream biological context as related but separable dimensions of microbiome interpretation.

Richness-oriented readouts were more sensitive to workflow transitions across Illumina and nanopore processing contexts. Much of this behavior was localized to rare-tail and low-abundance structure rather than to uniform restructuring of dominant taxa (Figure 1B and 1C; Supplementary Table 3). Workflow-associated richness differences also followed comparison-specific ratio and scaling behavior (Figure 2A to 2C). This interpretation is consistent with evidence that normalization and rarefaction choices can materially change microbiome results [19, 20, 15, 21]. Compositional constraints and differential-abundance method choice can also alter interpretation [23, 24, 25]. Rare or low-abundance features can be unstable or artifact-prone in amplicon data, with downstream effects on alpha diversity, community assembly, and network interpretation [26, 27].

The rare-tail findings do not imply that all rare taxa should be discarded. Low-abundance community members in the gut microbiome may carry ecological and host-relevant information [28]. A fixed abundance cutoff would therefore be too blunt for interpreting workflow-sensitive richness changes. The practical stance is to treat rare-tail-driven richness expansion as provisional until it persists beyond detection-boundary behavior. A signal becomes more robust when it remains after dominant-feature or cumulative-abundance restriction, consistent with the view that feature filtering and normalization should depend on the analysis goal rather than a universal cutoff [20, 46]. The same signal is further strengthened when it is visible in abundance-weighted summaries, because richness and abundance-weighted alpha-diversity measures can respond differently to rare features and sequencing depth decisions [19, 15, 21]. Greater caution is warranted when the signal is concentrated in low-abundance bins, becomes weaker after dominant features are analyzed separately, and is absent or weaker in abundance-weighted summaries. The present analysis examined low-abundance localization through abundance-bin and rank-abundance comparisons (Figure 1B and 1C). Rare-tail attenuation analyses then tested whether the signal weakened after dominant-only or cumulative-abundance restriction (Supplementary Table 3). Abundance-weighted comparisons assessed whether the same shift appeared beyond richness-based readouts (Figure 1A and Figure 2A). This operational approach preserves the biological possibility of low-abundance taxa while reducing the risk of overreading detection-sensitive features as robust ecological change.

The response-geometry framework builds on beta-space logic but focuses on paired response vectors rather than only on sample locations or group separation. Each participant was represented as a baseline-to-follow-up displacement in Aitchison CLR-Euclidean space (Figure 4B). This representation follows the broader logic of compositional data analysis, in which log-ratio geometry provides a principled coordinate system for relative-abundance data [31, 49, 22]. Response magnitude asks how far a participant moved. Directional coherence asks whether subject-level baseline-referenced response vectors are aligned within a defined feature space. Directional coherence is used here as an exploratory response-organization descriptor within that prespecified representation. It should not be treated as confirmatory unless it has a positive observed-minus-null gap, bootstrap or leave-one-out stability, reasonable zero-handling and feature-space sensitivity, endpoint alignment, and a prespecified analysis context. These conditions were not uniformly met here, so coherence is interpreted as an exploratory response-organization descriptor. Subject-level analyses then showed that displacement magnitude and directional alignment were related but not interchangeable (Figure 5A and 5B).

Pseudocount choice and zero handling can affect low-abundance features disproportionately in CLR-transformed microbiome data because zeros and near-zero counts are moved before log-ratio transformation [47, 48]. Zero replacement can therefore alter downstream log-ratio structure if it is not handled explicitly [47, 48]. Coherence can therefore vary with pseudocount choice. The present analyses therefore used a targeted pseudocount sensitivity check under fixed pairing, endpoint definitions, and feature filters. Changing the pseudocount from 0.5 to 1.0 did not materially alter the qualitative interpretation of the main synbiotic and dietary-intervention cohort. It also did not materially alter the qualitative interpretation of the FeFiFo 16S or endpoint-caveated FeFiFo CAZyme analyses. This check does not replace a systematic zero-handling benchmark, and the optional endpoint-matched CAZyme Fiber check was more sensitive. Weak or borderline coherence estimates should therefore not be read as stable evidence of coordinated response unless they are robust to zero handling, endpoint definition, and feature-space choice. The present study treats these estimates only as response-organization summaries within the specified feature space and endpoint definitions.

Simulation benchmarking was used as an internal operating check under simplified controlled assumptions before empirical intervention results were interpreted. Across 198,000 simulated datasets, null and magnitude-only random-direction scenarios showed observed-minus-null coherence gaps centered near zero. Random movement or large movement alone therefore did not generate coordinated directional organization (Supplementary Table 8; Supplementary Figures 6 and 8). Shared-direction simulations produced increasing coherence gaps and detection rates as effect size and sample size increased (Supplementary Table 8; Supplementary Figures 6 to 8). Mixed-responder simulations showed that pooled coherence can weaken when only a subset follows a shared direction (Supplementary Table 8; Supplementary Figures 6 to 8). Opposing-subgroup simulations showed that pooled coherence can cancel when subgroups move in opposing directions, even when displacement remains high (Supplementary Table 8; Supplementary Figures 6 to 8). Rare-tail simulations further supported the view that low-abundance feature inclusion can affect richness-sensitive summaries, while dominant-abundance attenuation reduces apparent richness change (Supplementary Table 8; Supplementary Figure 9). These simulations describe expected analytical behavior under the stated assumptions. They do not establish real-world calibration, universal false-positive control, clinical-use evidence, biological mechanism, or comparative performance against existing methods.

The 100-feature simulation setting is reported in the simulation parameter grid and operating summaries (Supplementary Table 8I). Its slightly liberal behavior is relevant for empirical interpretation because genus-level, pathway-level, or strongly filtered feature spaces can be similarly low-dimensional. Directional-coherence p values should therefore not be treated as stand-alone evidence of coordinated response in small feature spaces. Interpretation should also consider the observed-minus-null gap, bootstrap uncertainty, leave-one-out influence, pseudocount and zero-handling sensitivity, endpoint alignment, and the simulation operating context (Supplementary Table 8; Supplementary Figures 6 to 8). A nominally small p value alone is insufficient to claim coordinated response in low-dimensional feature spaces.

Supplementary implementation stress tests extended this operating check to empirical settings with stronger perturbation or large movement. The Palleja antibiotic-recovery trial-run outputs were used as a qualitative strong-perturbation implementation contrast only, based on the original antibiotic-recovery dataset ([37]; Supplementary Table 9). This contrast was not used to define an effect-size operating range for dietary-intervention applications. Antibiotic recovery, animal-diet shifts, and dietary or synbiotic remodeling differ in perturbation strength, temporal structure, baseline dependence, and feature-space geometry [36, 37]. The David animal-diet trial-run output was used as a magnitude-stress contrast, based on the original animal-diet intervention dataset ([36]; Supplementary Table 9). The available Animal-arm anchor transition showed large displacement without clear leave-one-out directional organization above the sign-flip null (Supplementary Table 9A). These analyses tested implementation behavior under specific public-data contrasts. They do not establish biological mechanism, clinical use, treatment effect, or general performance.

The synbiotic and dietary-intervention cohort can therefore be read as a practical example for separating response size from response organization in a modest intervention setting. The MetS-marker summaries are descriptive clinical-context information only. Paired microbiome displacement magnitude differed across intervention arms, but this pattern did not map simply onto MetS-marker redistribution (Figure 3A and 3B). DSG showed the clearest redistribution toward lower abnormal MetS-marker burden. After intervention, 17 of 18 DSG participants were in the 0MS or 1MS categories, compared with 15 of 18 in DG and 15 of 19 in SG. DSG also had the largest number of downward category shifts, with 8 of 18 participants shifting downward compared with 4 of 18 in DG and 3 of 19 in SG. The strongest microbiome displacement contrast was nevertheless observed between DG and DSG (Figure 3A and 3B; Supplementary Table 4). The methodological point is the mismatch between global taxonomic displacement and clinical-context redistribution. Global 16S-based compositional displacement summarizes community movement, but it cannot by itself represent intervention efficacy, between-arm clinical advantage, or physiological mechanism.

The direction-aware analyses further showed why magnitude alone is insufficient. Within-arm trajectories were heterogeneous, observed coherence remained close to the permutation-null reference, and bootstrap intervals broadly overlapped zero across DG, SG, and DSG (Figure 4B, 4C, and 4E). These results indicate limited pooled directional coherence in this cohort. Baseline microbiome differences may be large relative to the intervention effect. Similar exposure may therefore produce individualized trajectories rather than a single shared vector. Personalized diet-microbiome associations have been reported in human studies [1]. Individualized probiotic colonization responses have also been reported [6]. Baseline-dependent dietary-fiber response and individualized prebiotic or probiotic response potential have been reported in recent intervention and modeling studies [8, 9]. The simulation benchmark provides an operating reference for this interpretation. The closest available simulation grid to the cohort arm sizes was n = 20 rather than n = 18 or n = 19 (Supplementary Table 8G). The observed coherence summaries for the synbiotic and dietary-intervention cohort were small across arms, with matched observed-minus-null gaps ranging from −0.059 to −0.029 in the Figure 4 directional-coherence representation (Supplementary Table 8H). These values did not indicate coherence above the permutation-null reference. This simulation-referenced operating-context comparison supports interpreting the empirical pattern as limited pooled response organization rather than as a large shared-direction response.

Subject-level analyses showed that magnitude and directional coherence were partially related but not interchangeable. A weak positive rank association was observed between displacement magnitude and alignment to the group mean direction, but the group-adjusted linear model provided little support for a simple shared descriptor (Figure 5A). Directional alignment also varied broadly among subjects with similar displacement size (Figure 5B). Large microbiome movement can therefore arise from different response modes. Some subjects may move far in a direction shared with others, whereas others may move far toward distinct ecological states. An exploratory baseline-context analysis examined whether baseline microbiome features were associated with response magnitude or coherence (Supplementary Table 7D). This analysis remains secondary and hypothesis-generating. It is most useful as a rationale for future endpoint-aligned studies that evaluate baseline community state as a context variable for response magnitude or directional organization.

The simulation and stress-test results also help clarify apparently null or heterogeneous trial results. Absence of a large scalar diversity shift does not necessarily imply absence of microbiome response organization, because shared-direction simulations produced coherence increases even when interpretation depended on effect size and sample size (Supplementary Table 8; Supplementary Figures 6 to 8). A trial arm with low mean displacement but relatively high directional coherence could therefore reflect magnitude-limited but organized ecological remodeling. Large mean displacement with low directional coherence should not be interpreted as coherent intervention-associated response, because magnitude-only random-direction simulations remained near the null reference (Supplementary Table 8; Supplementary Figures 6 and 8). Participants may also move substantially but in ecologically divergent directions, as shown by opposing-subgroup simulations and the David animal-diet magnitude-stress contrast ([36]; Supplementary Table 8; Supplementary Table 9A). This distinction is especially relevant for dietary, fiber-focused, probiotic, and lifestyle microbiome interventions, where remodeling may be limited, distributed, heterogeneous, or individualized [1, 2, 6, 8]. Directional coherence is therefore best viewed as a supplementary exploratory descriptor that helps interpret scalar null or inconclusive results without converting them into efficacy claims.

The weak pooled organization observed in the synbiotic and dietary-intervention cohort motivated a second dietary-intervention application. The FeFiFo analysis asked whether the same response-organization logic could be applied across 16S rRNA gene amplicon taxonomic space and metagenomics-derived CAZyme gene-family feature space. The original FeFiFo study reported diet-specific multi-omic effects, including increased microbiota diversity after fermented-food exposure and increased microbiome-encoded glycan-degrading CAZymes after high-fiber exposure [2]. The present analysis did not re-test those primary outcomes. It added a response-organization layer to the existing taxonomic and functional-context data.

The FeFiFo results are interpreted here as layer-specific response-organization patterns. In the 16S rRNA gene amplicon analysis, the fermented-food arm showed more apparent taxonomic response organization than the high-fiber arm at the Baseline 0 to Week 14 endpoint (Figure 6D). The CAZyme analysis showed a small, endpoint-caveated high-fiber directional-coherence estimate in the retained metagenomics-derived GH/PL CAZyme RPM feature space (Figure 6E). This pattern is compatible with the diet-specific multi-omic context reported in the original FeFiFo study, but remains exploratory and feature-space dependent. Figure 6 therefore places adjacent response-organization applications in separate feature spaces. It should not be read as matched cross-layer confirmation.

Figure 6 should not be interpreted as a demonstration of matched multi-omic integration. The 16S rRNA gene amplicon and CAZyme analyses were placed in the same response-geometry framework because both can be represented as paired feature-space response vectors. Their endpoints were not matched, so taxonomic-functional coupling or concurrent cross-layer response cannot be inferred. The value of this comparison is methodological application across feature spaces, not matched multi-omics integration.

A matched-timepoint sensitivity check using the best-supported CAZyme-aligned endpoint, Baseline 0 to Week 8, further supported this boundary. Under this aligned endpoint and exact-subject 16S rRNA gene amplicon rerun, the endpoint-caveated Fiber-dominant CAZyme coherence pattern was not reproduced. This check reinforces that the FeFiFo CAZyme layer should be read as endpoint-sensitive, feature-space-dependent functional-context support (Supplementary Table 13C).

The CAZyme analysis adds a functional-context view of response organization, and its interpretation depends on endpoint timing and feature-space definition. The CAZyme endpoint was not identical to the primary 16S Week 14 endpoint. Exact primary-endpoint matching between 16S rRNA gene amplicon and CAZyme profiles was not available in the processed objects used here (Supplementary Tables 10 and 13). No primary-endpoint cross-layer inference was attempted. The retained feature space covered 840 metagenomics-derived GH/PL CAZyme RPM feature IDs, comprising GH and PL family-plus-subfamily features in the predefined functional-context analysis (Figure 6E; Supplementary Tables 12 and 13). This space was broader than the 11 CAZymes reported as increased in the original high-fiber analysis [2]. The original upregulation analysis identifies specific CAZyme-annotated features that increased in abundance. Directional coherence asks whether participant-level CAZyme profile changes moved in a similar multivariate direction. Its scope is multivariate profile organization rather than catalytic function, metabolic flux, mechanism, or matched-primary-endpoint parallelism with the 16S pattern. The CAZyme result is also feature-space dependent. Feature-retention sensitivity support showed that the current retained GH/PL feature space and a stricter prevalence rule retained the same 840 features (Supplementary Table 13B). Top-abundance and top-variance subset checks supported feature-space dependence (Supplementary Table 13B). A substantially different CAZyme retention rule or zero-handling strategy could therefore change the high-fiber estimate.

This positioning clarifies how the framework relates to existing methods. The framework does not replace PERMANOVA, PERMDISP, RPCA, longitudinal microbiome workflows, or longitudinal log-ratio approaches. PERMANOVA tests group-location or centroid structure [16]. PERMDISP tests dispersion structure, and multivariate dispersion can also be interpreted as a beta-diversity descriptor [17, 3]. RPCA and other compositional ordination approaches provide important tools for representing sparse microbiome perturbations [32]. Longitudinal microbiome workflows support paired and repeated-measures analysis [29]. Repeated-measures visualization and log-ratio approaches can further improve interpretation of temporal microbiome data [35, 30]. Directional coherence asks a different paired-response question. It asks whether subject-level baseline-referenced response vectors are aligned after displacement from baseline has been quantified. These questions can diverge in heterogeneous response settings. Directional coherence should therefore be used as a complementary response-organization descriptor, not as a replacement analysis.

Several limitations define how these findings should be interpreted. Statistical precision remains the main constraint for the synbiotic and dietary-intervention cohort. The arm sizes were modest, which limited the ability to distinguish weak directional structure from background heterogeneity or to assess weaker and subgroup-specific organization. This constraint was reflected by broad bootstrap uncertainty around coherence estimates (Figure 4E). Bootstrap intervals and leave-one-out checks provide stability diagnostics within the observed cohort, but they do not support external generalization.

Technical attribution is another constraint. Workflow and platform comparisons were compound analytical-context comparisons, and sequencing platform, reference framework, clustering behavior, zero handling, and pipeline versions were not fully separable in every contrast. Systematic zero-handling benchmarking remains incomplete. The targeted 0.5 versus 1.0 pseudocount sensitivity check did not materially alter the main qualitative calls for the synbiotic and dietary-intervention cohort, FeFiFo 16S, or endpoint-caveated FeFiFo CAZyme analyses (Supplementary Table 5; Supplementary Table 13). These analyses support caution about analytical sensitivity, but they do not attribute each observed shift to a single technical source.

Method generalization also remains limited. The coherence statistic was used as a descriptive and robustness-supported response-organization summary. The r500 benchmark provides an internal operating check under simplified controlled assumptions (Supplementary Table 8; Supplementary Figures 6 to 9). It did not fit empirical covariance matrices, cohort-specific temporal dependence, empirical sequencing-depth structure, or the full taxon-correlation structure of real intervention datasets. It does not establish real-world calibration, universal false-positive control, clinical-use evidence, biological mechanism, or comparative performance against existing methods. The present analysis therefore interprets coherence relative to the permutation-null, bootstrap uncertainty, sensitivity checks, endpoint alignment, and simulation context, rather than by applying a fixed cutoff. The David/Palleja stress tests provide supplementary empirical operating contrasts under strong-perturbation and magnitude-stress settings (Supplementary Table 9).

The clinical and functional-context analyses were designed as exploratory applications rather than primary endpoints, reflecting the framework’s current stage of development. The synbiotic and dietary-intervention cohort clinical-context summaries should not be used to infer efficacy, between-arm clinical advantage, or participant classification. The FeFiFo CAZyme analysis was constrained by endpoint mismatch and by the fact that CAZyme RPM profiles represent metagenomics-derived gene-family abundance, not catalytic function or direct metabolic flux (Supplementary Tables 10 to 14).

Future studies should align taxonomic, functional, and host-context measurements at the same endpoints and follow contemporary microbiome-study reporting principles emphasizing transparent workflow description, metadata reporting, and data/code availability [51]. They should also prespecify response-organization analyses, compare zero-handling strategies, and evaluate whether baseline community state is associated with response magnitude or directional organization in external datasets with interpretable host-context labels. Temporal community methods could be useful complements because they provide established ways to analyze time-dependent community change [33, 34]. Longitudinal microbiome design tools could also help organize paired or repeated measurements before response-vector summaries are interpreted [29]. Such designs would allow directional coherence to be assessed as a prespecified exploratory descriptor rather than as an inferential endpoint. To support transparent reuse, the accompanying beta research-preview GitHub repository provides documented scripts, toy examples, environment notes, output-interpretation guidance, pseudocount sensitivity utilities, a simulation-based operating guide, and a script-based exploratory implementation of the response-geometry workflow. The repository is intended to support reproducibility inspection and exploratory reuse, not clinical, diagnostic, predictive, efficacy, regulatory, production, or mechanistic inference. The repository is intended to support reproducibility inspection and exploratory reuse, not clinical, diagnostic, predictive, efficacy, or mechanistic inference.

## Conclusions

This study presents a conservative magnitude-coherence framework for interpreting microbiome remodeling during intervention periods, especially in mild and heterogeneous intervention settings. Its core contribution is to separate analytical sensitivity, baseline-referenced displacement, and participant-level directional organization. Figure 7 summarizes this decision logic and keeps downstream biological context as a related but separate interpretive layer.

The framework is a proof-of-concept response-organization descriptor, not an established inferential endpoint. It can describe when taxonomic or functional-context profiles show coordinated movement within a defined feature space. It also limits the risk of treating rare-tail-sensitive richness shifts, magnitude-only displacement, or heterogeneous movement as shared ecological response. Future endpoint-aligned intervention studies could evaluate whether directional coherence is useful as a prespecified exploratory descriptor alongside diversity, displacement magnitude, PERMANOVA/PERMDISP, and beta-diversity summaries. This use would help distinguish organized response structure from heterogeneous movement without converting weak, null, or endpoint-caveated findings into efficacy or mechanistic claims.

## Supporting information

List of supplemental files

Supplementary Table 1 - Numerical support for Figure 1A-C

Supplementary Table 2 - Statistical support for Figure 2 comparison and scaling summaries

Supplementary Table 3 - Attenuation-based support for lower-abundance concentration of workflow-associated observed-richness contrasts

Supplementary Table 4 - Statistical support for Figure 3B microbiome displacement magnitude comparisons

Supplementary Table 5 - Directional-coherence support summary for Figure 4

Supplementary Table 6 - Divergence support summary for Figure 4

Supplementary Table 7 - Exploratory baseline predictor support for Figure 5

Supplementary Table 8 - r500 simulation benchmarking tables for the response-geometry framework.

Supplementary Table 9 - David/Palleja strong-perturbation implementation stress tests

Supplementary Table 10 - Wastyk external dataset and endpoint summary.

Supplementary Table 11 - Wastyk 16S framework results.

Supplementary Table 12 - Wastyk CAZyme functional-coherence results.

Supplementary Table 13 - Wastyk 16S-versus-CAZyme response-space comparison.

Supplementary Table 14 - Wastyk evidence hierarchy and host-context caveat summary.

## List of abbreviations

ASV: amplicon sequence variant
CAZyme: carbohydrate-active enzyme
CLR: centered log-ratio
DG: dietary intervention group
DSG: combined dietary plus synbiotic group
FeFiFo: Fermented and Fiber-rich Foods
GH: glycoside hydrolase
MetS: metabolic syndrome
OTU: operational taxonomic unit
PCoA: principal coordinates analysis
PERMANOVA: permutational multivariate analysis of variance
PERMDISP: permutational analysis of multivariate dispersions
PL: polysaccharide lyase
RPM: reads per million
SG: synbiotic supplementation group

## Declarations

### Ethics approval and consent to participate

This study reanalyzed publicly available and previously published de-identified datasets. Ethics approval and participant consent for the original studies were obtained by the original investigators as described in the cited source publications. No new human participants, human tissue, animal experiments, or prospective data collection were included in this secondary methodological analysis.

### Consent for publication

Not applicable.

### Availability of data and materials

The datasets analysed during the current study are publicly available in the repositories and accession records described in the Methods section and cited source publications. Raw public datasets are not redistributed in the accompanying repository. A beta research-preview GitHub repository accompanying the preprint is available at https://github.com/carolyyszeto/microbiome-response-interpreter-beta and has been tagged as v6.5-beta. The repository provides documented scripts, a toy dataset, environment notes, output-interpretation guidance, pseudocount sensitivity utilities, a simulation-based operating guide, and exploratory implementation support for the response-geometry workflow. This repository is intended as a preprint companion and reproducibility-inspection resource rather than a validated clinical, diagnostic, predictive, efficacy, regulatory, production, or mechanistic tool.

### Competing interests

The authors declare that they have no competing interests.

### Funding

No funding was received for this study.

### Authors’ contributions

CYYS conceived the study, performed the analyses, interpreted the results, and drafted the manuscript. HSK reviewed the manuscript. All authors read and approved the final manuscript.

## Acknowledgements

Not applicable.

## Authors’ information

Not applicable.

## Notes

### Competing Interest Statement

The authors have declared no competing interest.

