## Supplementary figures 1-11 for "A response-geometry framework separates microbiome movement magnitude from directional coherence in intervention studies"

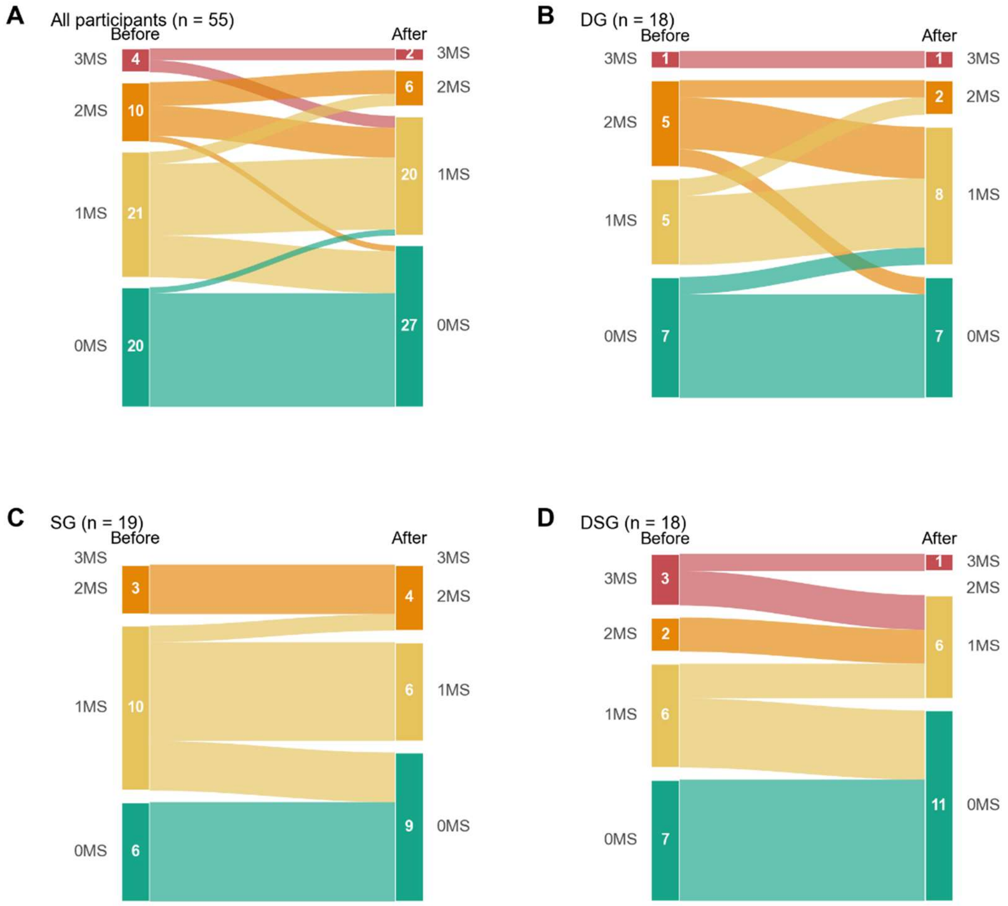


Supplementary Figure 1. Transition of abnormal MetS-marker count categories across intervention groups. (A) All paired participants (`n = 55`), (B) DG (`n = 18`), (C) SG (`n = 19`), and (D) DSG (`n = 18`). Alluvial ribbons connect before- and after-intervention abnormal MetS-marker count categories, where `0MS`, `1MS`, `2MS`, and `3MS` denote the presence of `0`, `1`, `2`, or `3` abnormal MetS-related markers among fasting glucose, HDL-C, and triglycerides. Ribbon width is proportional to participant counts, and ribbon color encodes the before-intervention category. This supplementary figure extends the participant-level transition framing underlying main Figure 3A. Formal statistical support for the microbiome displacement panel in main Figure 3B is provided separately in `figure3_panelB_stats_table.csv`.

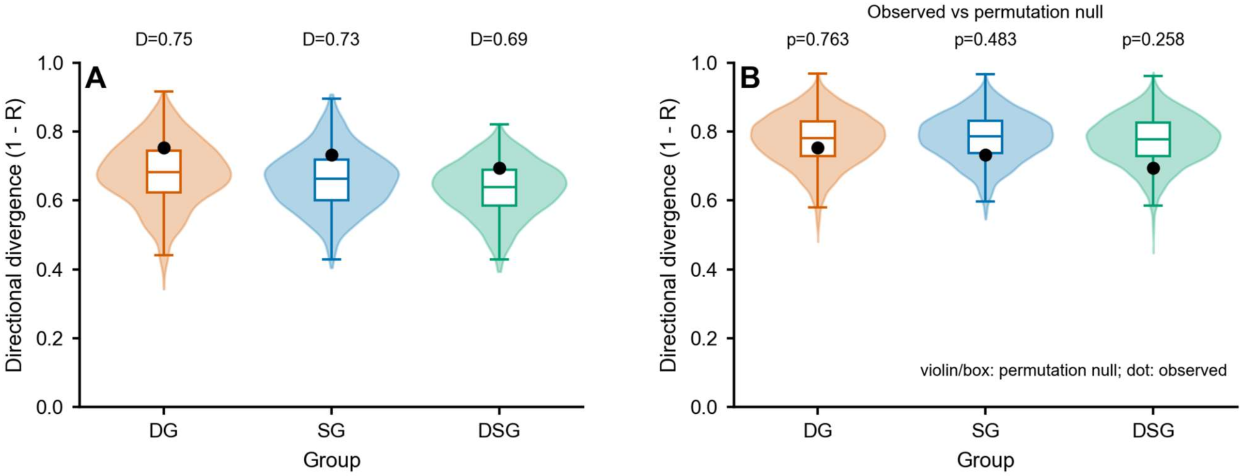


Supplementary Figure 2. Directional divergence support for Figure 4C–D. (A) Observed directional divergence, defined as $D = 1 - R$, across DG, SG, and DSG in the $K = 5$ Aitchison CLR-Euclidean representation. Larger values indicate greater heterogeneity of response direction within the group. (B) Observed divergence compared with a within-subject sign-flip permutation null. Across groups, observed divergence remained close to the null reference, supporting the interpretation that directional heterogeneity was present but not clearly separated from null expectation in the current dataset. This figure is retained as secondary support for the heterogeneity reading relevant to Figure 4C–D and should not displace the coherence-first supplementary interpretation.

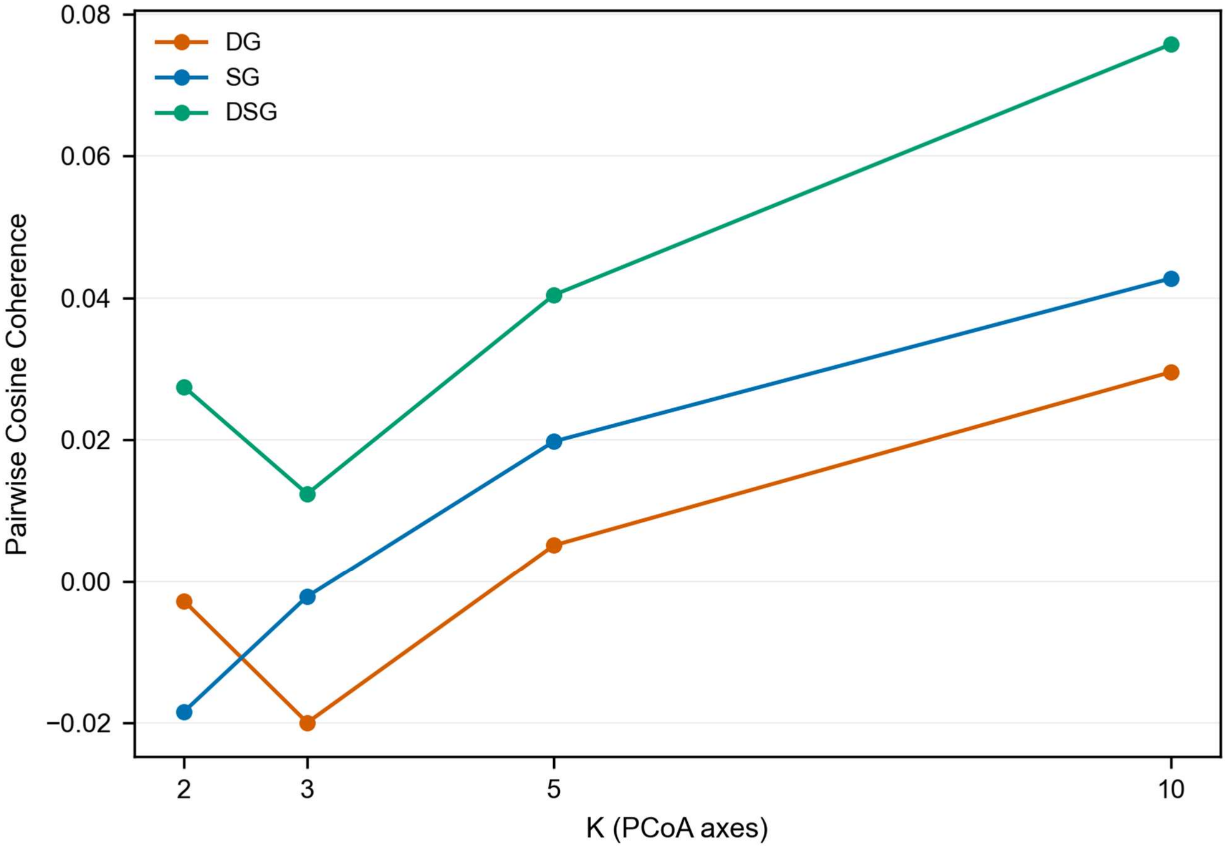


Supplementary Figure 3. K-sensitivity coherence support for Figure 4C. This figure shows the sensitivity of the directional-coherence statistic to the number of retained CLR-Euclidean axes (K) used to define paired response vectors. Across DG, SG, and DSG, coherence estimates remained low across the tested K settings, indicating that the main null-leaning interpretation of weak within-arm directional coordination was not driven by a single arbitrary choice of ordination dimensionality. This figure is presented as robustness support rather than as a separate lead result.

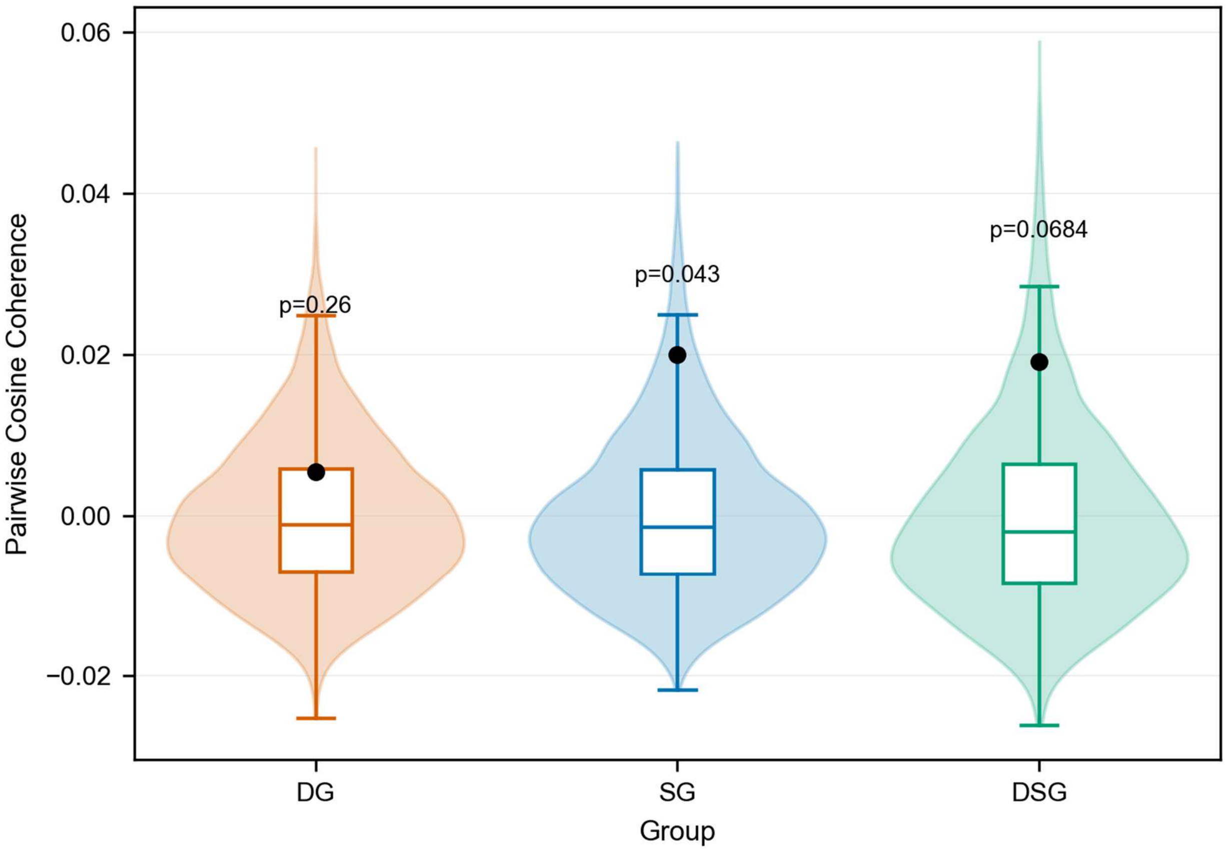


Supplementary Figure 4. CLR-space permutation sensitivity support for Figure 4C. This figure evaluates directional coherence in full CLR space using a permutation-based null that preserves within-subject pairing while disrupting consistent directional alignment. Across DG, SG, and DSG, observed coherence remained close to the null reference, supporting the interpretation that within-arm directional coordination was weak in the current dataset. This figure is intended as a sensitivity analysis supporting the Figure 4 coherence reading.

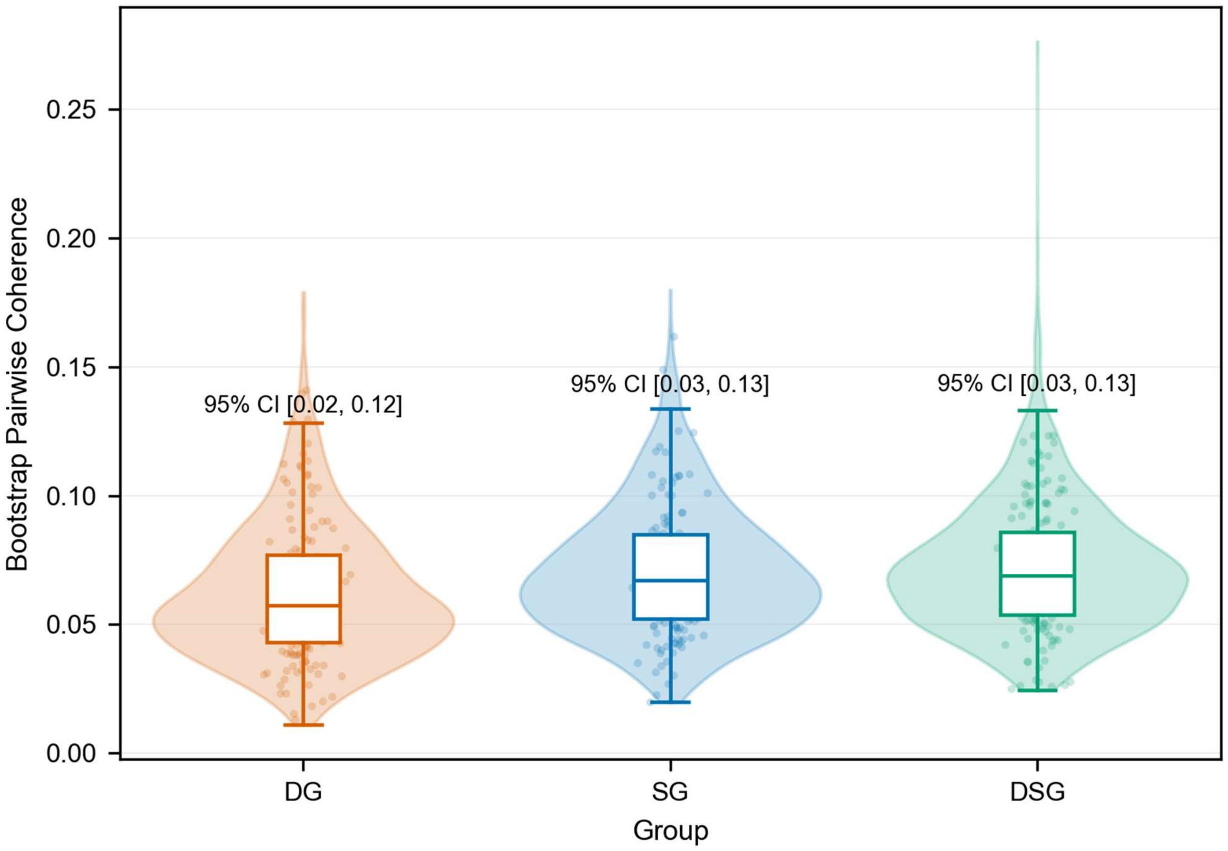


Supplementary Figure 5. CLR-space bootstrap robustness support for Figure 4E. This figure shows bootstrap resampling distributions for the directional-coherence statistic in full CLR space across DG, SG, and DSG. Confidence intervals broadly overlapped zero across groups, indicating that the coherence estimates were weak and uncertain under resampling. This bootstrap view supports the main conclusion that strong within-arm directional organization was not established in the current dataset.

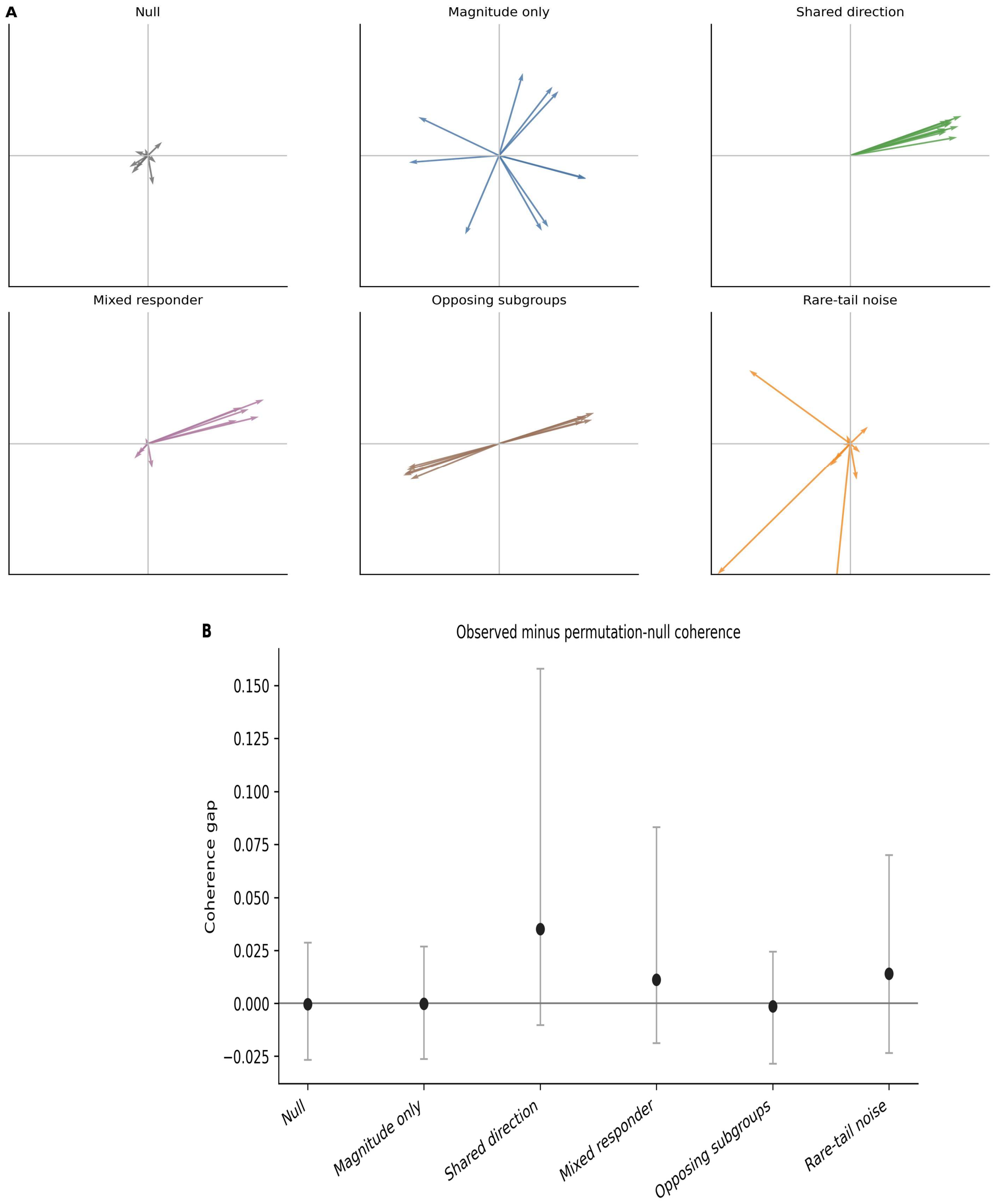


Supplementary Figure 6. Simulation scenarios and coherence-gap behavior. (A) shows simulated response scenarios, including null response, magnitude-only random-direction response, shared-direction response, mixed-responder response, opposing responder subgroups, and rare-tail noise. Arrows represent participant-level baseline-to-follow-up response vectors in CLR-Euclidean space. (B) shows observed-minus-null coherence gap by scenario, defined as observed group coherence minus the permutation-null mean. Null and magnitude-only scenarios remained near zero, shared-direction simulations showed the strongest positive gap, mixed-responder simulations showed attenuated coherence, and opposing-subgroup simulations showed low pooled coherence. These simulations evaluate expected analytical behavior under controlled assumptions and do not prove biological mechanism or clinical response.

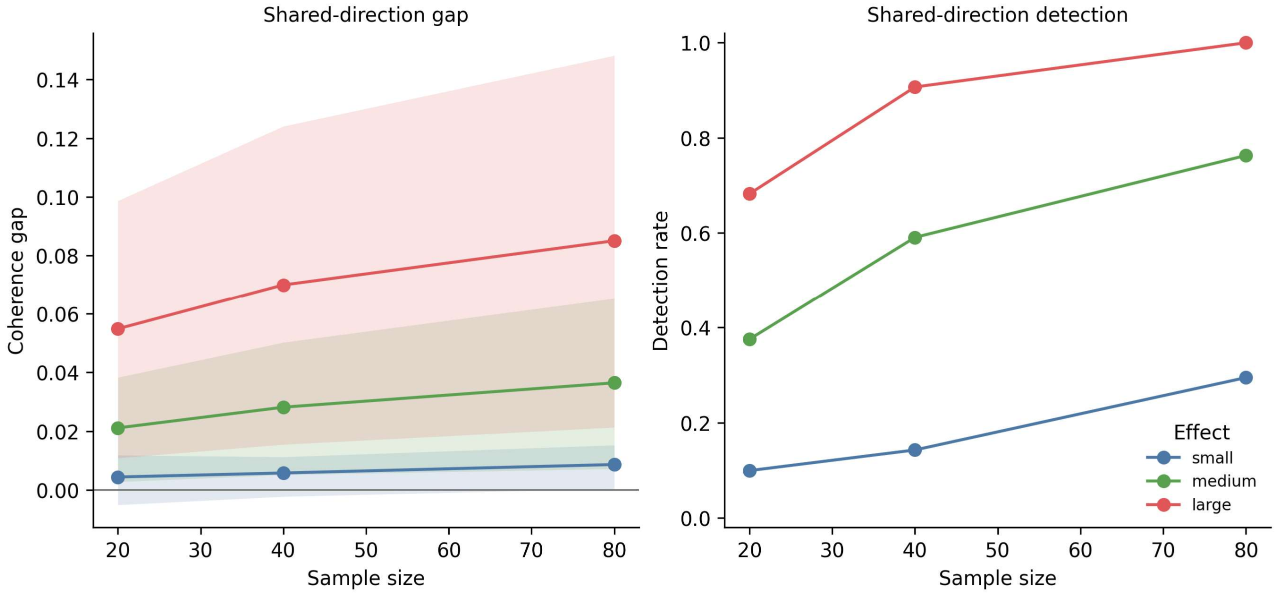


Supplementary Figure 7. Shared-direction positive-control behavior. Shared-direction simulations showed increasing coherence gap and detection rate with larger effect size and sample size. This positive-control scenario evaluates whether the response-geometry framework detects coordinated response direction when a common simulated direction is imposed. Detection behavior is interpreted as diagnostic support for expected analytical behavior, not as full calibration of a formal hypothesis test.

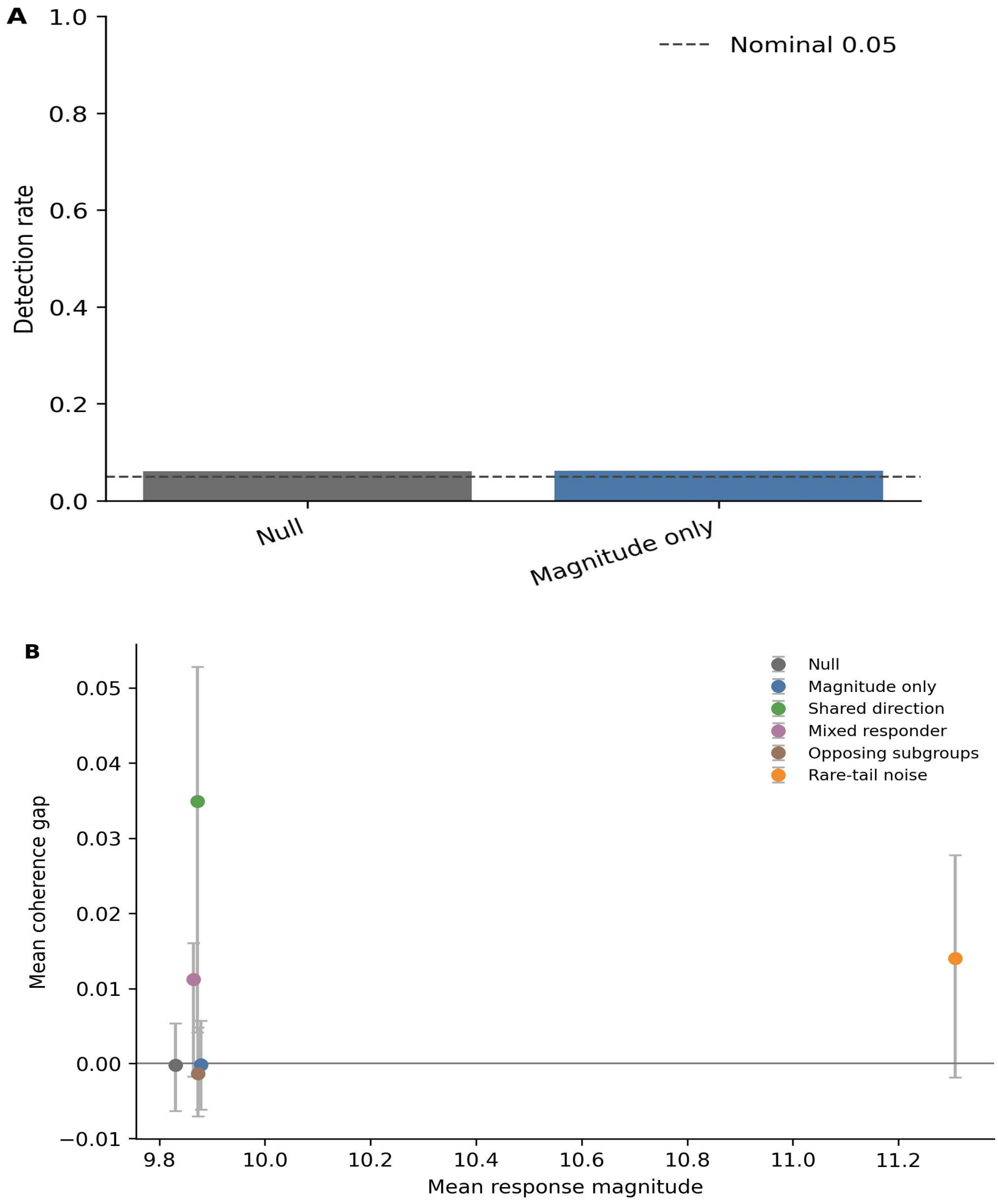


Supplementary Figure 8. False-positive reference behavior and magnitude-coherence separation. (A) shows empirical detection under null and magnitude-only random-direction scenarios. Detection was close to the nominal reference but slightly liberal in the aggregate simulation grid, especially in lower-dimensional feature spaces. (B) shows that response magnitude and coherence gap are distinct quantities; magnitude-only and opposing-subgroup scenarios can show substantial displacement without strong pooled directional organization. Coherence should be interpreted as a response-organization summary rather than a standalone calibrated inferential endpoint.

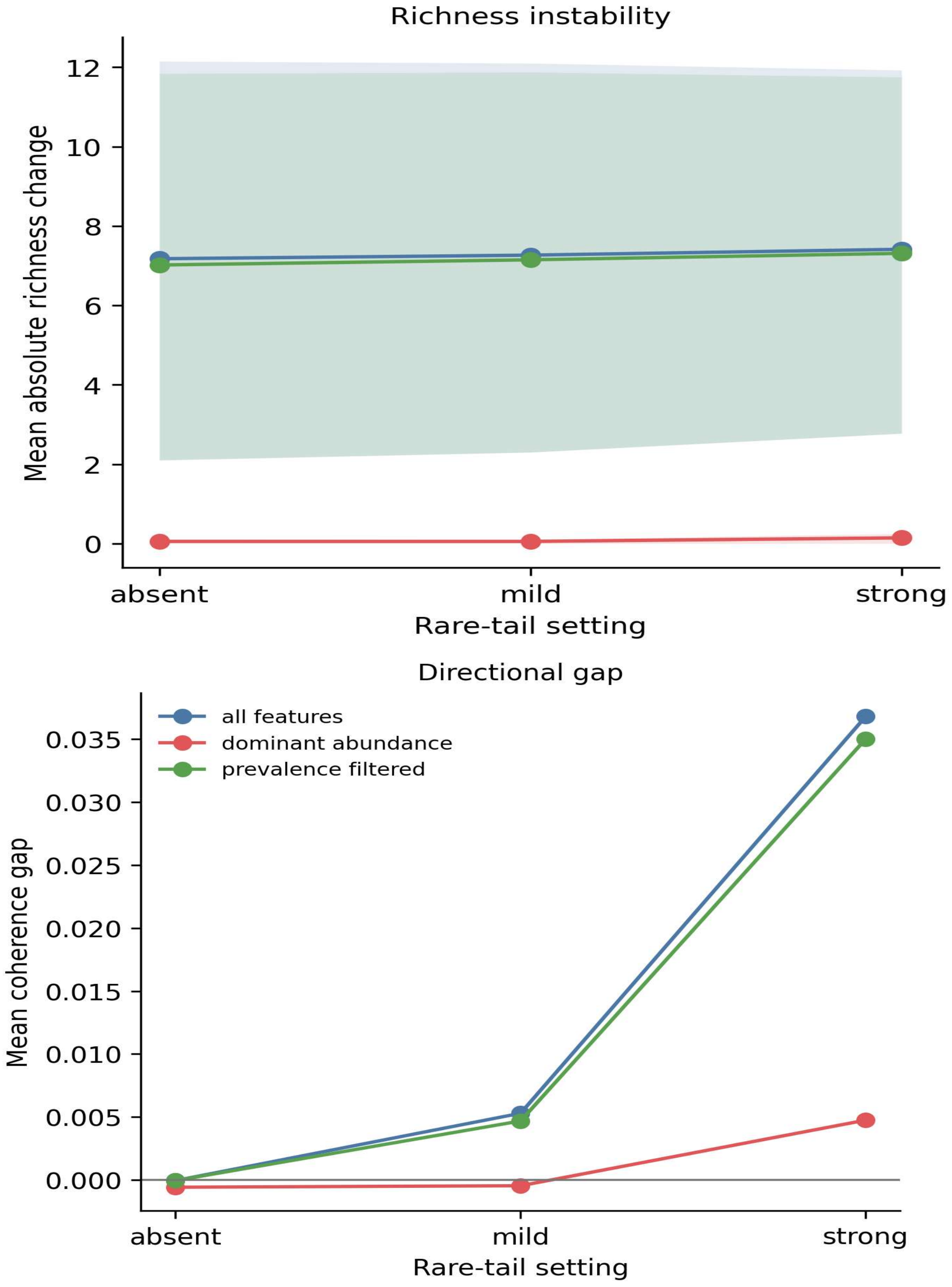


Supplementary Figure 9. Rare-tail attenuation reduces richness-sensitive change. Rare-tail sensitivity analyses compared all-feature, prevalence-filtered, and dominant-abundance feature sets. Low-abundance-retaining analyses showed large richness-sensitive changes, whereas dominant-abundance attenuation strongly reduced the apparent richness shift. The rare-tail result supports feature-inclusion sensitivity rather than a clean monotonic rare-tail-strength dose response.

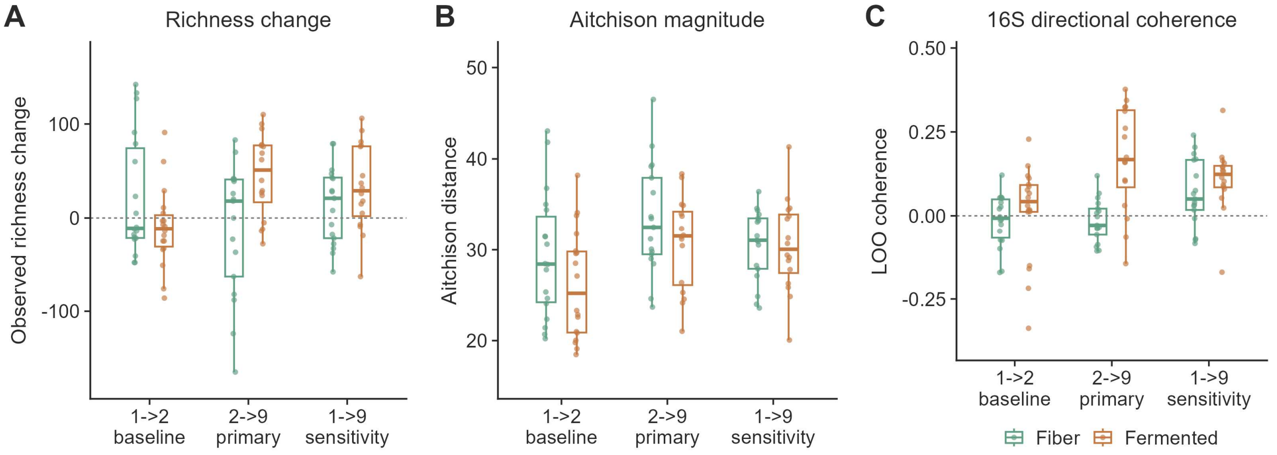


Supplementary Figure 10. Wastyk 16S external-application support. (A) Subject-level observed-richness change across the 1→2 baseline interval, the 2→9 primary 16S endpoint, and the 1→9 sensitivity contrast in the high-fiber and fermented-food arms. (B) Subject-level Aitchison displacement magnitude across the same contrasts. (C) Subject-level leave-one-out directional coherence across the same contrasts, measured as cosine similarity to the group-mean response direction estimated without the focal subject. Points denote paired participants, boxes show interquartile ranges, center lines show medians, and dashed horizontal lines indicate no change or zero coherence where applicable. The 2→9 contrast represents the primary 16S rRNA gene amplicon endpoint, whereas 1→9 is shown as a sensitivity contrast.

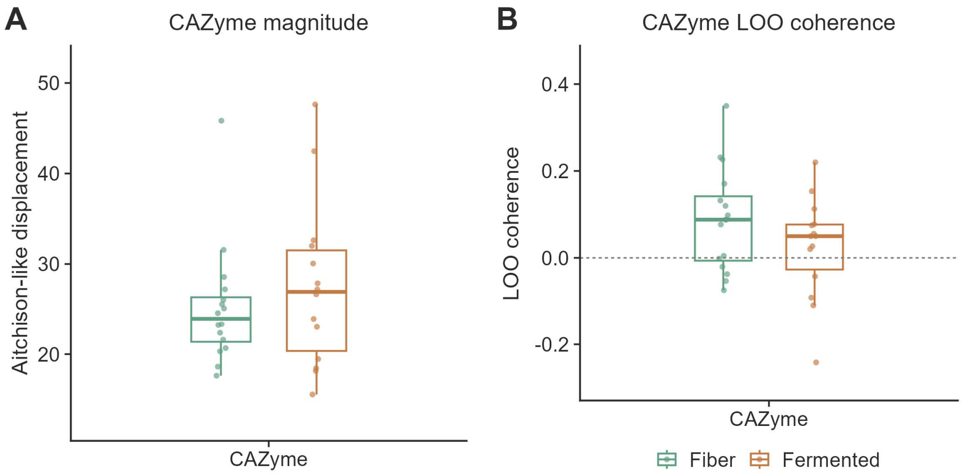


Supplementary Figure 11. Wastyk CAZyme functional-coherence support. (A) CAZyme feature-space displacement magnitude in the high-fiber and fermented-food arms. (B) CAZyme leave-one-out directional coherence in the same arms, measured as cosine similarity to the group-mean CAZyme response direction estimated without the focal subject. Points denote paired participants, boxes show interquartile ranges, center lines show medians, and the dashed horizontal line indicates zero coherence. CAZyme profiles were metagenomics-derived and analyzed using the available baseline-to-end-maintenance contrast; this endpoint is not identical to the primary 16S Week 14 endpoint and is therefore interpreted as endpoint-caveated functional context.
